# Canonical TGFβ signaling induces collective invasion in colorectal carcinogenesis through a Snail1- and Zeb1-independent partial EMT

**DOI:** 10.1101/2020.12.11.420851

**Authors:** Marion Flum, Severin Dicks, Monika Schrempp, Alexander Nyström, Melanie Boerries, Andreas Hecht

## Abstract

Local invasion is the initial step towards metastasis, the main cause of cancer mortality. In human colorectal cancer (CRC), malignant cells predominantly invade as cohesive collectives, and may undergo partial epithelial-mesenchymal transition (pEMT) at the invasive front. How this particular mode of stromal infiltration is generated is unknown. Here we investigated the impact of oncogenic transformation and the microenvironment on tumor cell invasion using genetically engineered organoids as CRC models. We found that inactivation of the *Apc* tumor suppressor combined with expression of oncogenic *Kras^G12D^* and dominant negative *Trp53^R172H^* did not cell-autonomously induce invasion *in vitro*. However, oncogenic transformation primed organoids for activation of a collective invasion program upon exposure to the prototypical microenvironmental factor TGFβ1. Execution of this program co-depended on a permissive extracellular matrix which was further actively remodeled by invading organoids. Although organoids shed some epithelial properties particularly at the invasive edge, TGFβ1-stimulated organoids largely maintained epithelial gene expression while additionally implementing a mesenchymal transcription pattern, resulting in a pEMT phenotype that did not progress to a fully mesenchymal state. Induction of this stable pEMT required canonical, Smad4-mediated TGFβ signaling, whereas the EMT master regulators Snail1 and Zeb1 were dispensable. Gene expression profiling provided further evidence for pEMT of TGFβ1-treated organoids and showed that their transcriptomes resemble those of human poor prognosis CMS4 cancers which likewise exhibit pEMT features. We propose that collective invasion in colorectal carcinogenesis is triggered by microenvironmental stimuli through activation of a novel, transcription-mediated form of non-progressive pEMT independently of classical EMT regulators.

## Introduction

Metastasis - accountable for the overwhelming majority of cancer-related deaths in solid cancers - requires that tumor cells successfully pass through a series of events summarily termed invasion-metastasis cascade (1). Tumor cells can accomplish the initial step of this cascade by shedding cell-cell contacts and infiltrate adjacent stromal tissue as individual cells, employing amoeboid or mesenchymal modes of migration (2). However, most human cancers display a collective mode of invasion where tumor cells maintain intercellular interactions and migrate in cohesive groups (2). Nonetheless, cells at the invasive front of such a collective may also display mesenchymal features (2, 3). Unfortunately, knowledge about the prerequisites and molecular determinants which promote a specific mode of invasion is limited.

Epithelial-mesenchymal transitions (EMT) are complex cellular programs that were repeatedly implicated in cancer cell invasion and metastasis (4). In the course of EMT, cells gradually trade key epithelial characteristics such as apical-basal polarity, and tight cell-cell and cell-matrix contacts for mesenchymal features including fibroblast-like morphology, front-rear polarity, increased motility, and enhanced invasiveness. Cancer cells can be induced to undergo EMT in response to extrinsic and intrinsic stimuli, for example from the Wnt, Notch, mitogen-activated protein kinase (MAPK), and TGFβ signaling pathways (4). Irrespective of the upstream trigger, a central role in EMT processes is typically attributed to a small group of transcription factors (TFs) from the Snail, Zeb, and Twist families (5). These EMT-TFs are thought to orchestrate the transition from epithelial to mesenchymal states by extensive transcriptional reprogramming. Yet, recent studies suggest that cancer cells may not need to acquire a fully mesenchymal phenotype in order to attain maximum invasive and metastatic capacity. Rather, the ability of cancer cells to traverse only partway through EMT and to present with variable combinations of epithelial and mesenchymal properties appears to be most advantageous for metastasis (6, 7). Although instances of such partial EMT (pEMT) could be captured *in vitro* and *in vivo* (6, 8–12), it is not clear whether pEMT states only represent snapshots along a continuum of intermediate states towards complete EMT (cEMT) (8, 12), or endpoints of independent pEMT programs. Furthermore, it also became evident that tissue-specific and transcription-independent variants of pEMT exist (9, 10).

Colorectal cancer (CRC) is one of the most frequent forms of cancer worldwide. Histological examination indicated that the predominant form of stromal infiltration in CRC is collective invasion with evidence for pEMT at the invasive front (3). How this particular pattern of invasiveness arises is unknown. Work with genetically engineered mouse models and organoids suggested that the three most common tumor-promoting events in CRC, disruption of the *Apc* and *Trp53* tumor suppressor genes in conjunction with oncogenic mutations in *Kras*, suffice to induce invasion and metastatic disease (13–16), but it was not determined whether the experimental models recapitulated the particular type of invasion observed in CRC tissue specimens. Furthermore, there are contradictory results concerning the number and type of genetic changes needed to elicit invasion and metastasis (15, 17–20), and an invasion-stimulating effect of the tumor microenvironment (TME) cannot be excluded.

TGFβ family members are prototypic examples for microenvironmental factors with the capacity to induce cancer cell invasion. However, the role of TGFβ signaling in CRC is quite controversial. TGFβ receptors and Smad4, a key TF in canonical TGFβ signaling, are frequently inactivated to boost experimental metastasis in animal models (15, 17–21), and it was proposed that TGFβ signaling might act on nonneoplastic rather than cancer cells to promote metastases formation (22, 23). Yet, the TGFβ pathway appears mostly functional in CRC specimens (24), and comprehensive transcriptome analyses provided evidence for active TGFβ signaling in the consensus molecular subtype 4 (CMS4) of human CRC, which is the CMS with the poorest prognosis (25). Likewise, TGFβ pathway activity was documented in a metastatic mouse tumor model (13), and TGFβ signaling induced invasion and installed CMS4-like transcription in a model of the sessile serrated adenoma CRC subtype (26). Even though TGFβ signaling typically induces cEMT and single cell invasion (8, 10, 12, 27), it therefore could be that invasive behavior in CRC is dual a consequence of cell-autonomous and non-autonomous mechanisms.

Here, we aimed to dissect the impact of tumor cell genetics and extrinsic factors on invasive behavior in intestinal tumorigenesis by employing genetically modified murine organoids under defined conditions. We report that disruption of *Apc* and *Smad4* together with expression of Kras^G12D^ and the dominant negative p53^R172H^ mutant do not suffice to elicit cell-intrinsic invasiveness. However, concomitant oncogenic lesions in the Wnt, MAPK, and p53 signaling pathways primed organoids for TGFβ1-inducible collective invasion and a stable, transcription-dependent pEMT which was executed independently of Snail1 and Zeb1. Interestingly, TGFβ1-treated organoids acquired a gene expression pattern representative of CMS4 suggesting that an atypical TGFβ1 response and a novel pEMT variant may underlie human CRC collective invasion.

## Results

### Intestinal organoids do not gain cell-intrinsic invasiveness by oncogenic transformation

To test whether hyperactive Wnt and MAPK signaling and impaired p53 activity induce invasiveness in a cell-autonomous fashion, we generated multiple small intestinal organoid lines from *Apc^580S/580S^*, *Kras^LSL-G12D/+^*, *Trp53^LSL-R172H/+^*, *tgVillinCreER^T2^* mice (Fig. 1a, Supplementary fig. 1). Organoids (hereafter termed floxed organoids) were treated with 4-hydroxy-tamoxifen (4-OHT) *in vitro* to obtain *Apc*-deficient organoids expressing oncogenic Kras^G12D^ and dominant negative p53^R172H^ (hereafter called TKA organoids; Supplementary fig. 1a-d). Floxed and TKA organoids were analyzed for evidence of oncogenic transformation and invasiveness. As observed before (17, 18), TKA organoids exhibited cystic growth when compared to bud-forming floxed organoids (Fig. 1b) and remained viable in media without R-spondin-1 (Supplementary fig. 1e, f). TKA organoids were also EGF-independent and tolerated EGFR inhibition (Supplementary fig. 1e, f). Furthermore, expression of Wnt target genes and intestinal stem cell markers increased in TKA organoids while that of differentiation markers decreased (Supplementary fig. 1g). This suggests an expansion of stem and progenitor cells in TKA organoids at the expense of multilineage differentiation. Notably, acquired growth factor independence and disturbed differentiation represent hallmarks of cancer cells. Yet, TKA organoids remained non-invasive. Like floxed organoids, they were surrounded by a continuous layer of laminin and showed apical localization of atypical protein kinase C (aPKC) (Fig. 1c). Forskolin-inducible swelling of organoids confirmed epithelial integrity (28), albeit floxed and TKA organoids displayed different swelling dynamics which might be caused by the differences in shape and elasticity of organoids (Fig. 1d). Thus, TKA organoids maintained apico-basolateral polarity, basement membrane integrity, and functional cell-cell junctions which are key characteristics of epithelial cell layers. There was also no indication for invasiveness when organoids were cultivated in an air-liquid interface setup with type I collagen to expose them to an extracellular matrix (ECM) more representative of a desmoplastic tumor microenvironment (29). The majority of non-transformed floxed organoids perished under these conditions, whereas TKA organoids remained viable and occasionally showed evidence of dysplasia (Fig. 1e). However, TKA organoids did not infiltrate the type I collagen matrix. In summary, despite several lines of evidence for oncogenic transformation, TKA organoids did not display cell-intrinsic invasiveness arguing that invasive behavior observed *in vivo* might be triggered by cell non-autonomous mechanisms.

**Fig. 1:**
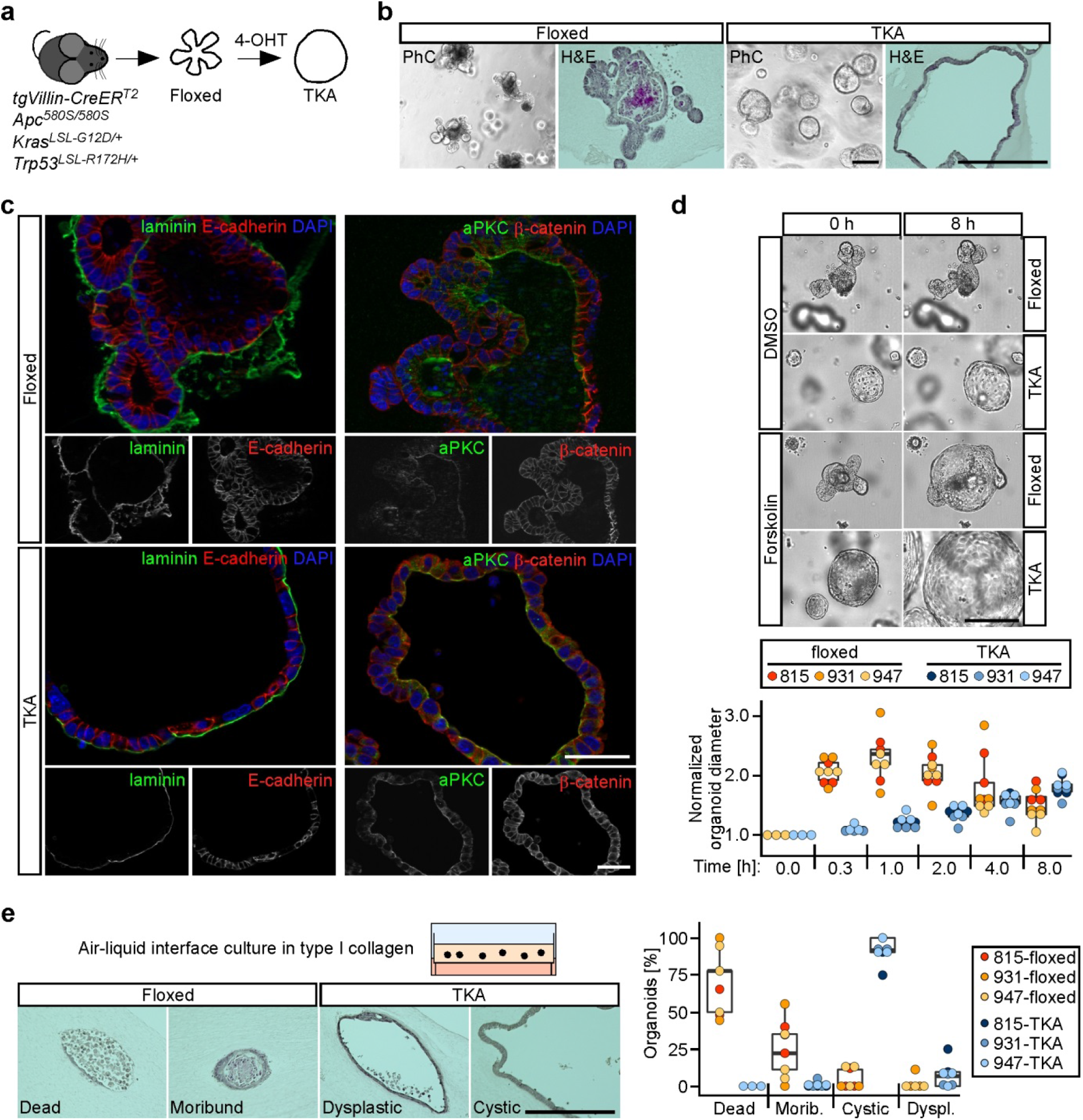
Oncogenically transformed intestinal organoids display no cell-intrinsic invasiveness *in vitro*. **a**, Strategy for generating oncogenically transformed small intestinal and colonic organoids from genetically engineered mice carrying a *Villin-CreER^T2^* transgene, two floxed *Apc* alleles, as well as heterozygous *Kras^LSL-G12D/+^* and *Trp53^LSL-R172H/+^* loci. Floxed stop cassettes (LSL) prevent expression of the mutant *Kras* and *Trp53* alleles. For recombination, organoids were treated with 4-hydroxy-tamoxifen (4-OHT), yielding TKA organoids. **b**, Whole mounts and hematoxylin/eosin (H&E) stained sections of floxed and TKA organoids (line 815) cultured in 7 mg/ml Matrigel and visualized by phase contrast (PhC) or bright field microscopy. Scale bars: 200 μm. **c**, Representative images of immunofluorescence staining of pan-laminin, E-cadherin, atypical protein kinase C (aPKC), and β-catenin in sections of floxed and TKA organoids (line 815) cultured in 7 mg/ml Matrigel. Nuclei were stained by DAPI; n>3. Scale bars: 50 μm. **d,** Top: Microscopy of floxed and TKA organoids (line 815) in 7 mg/ml Matrigel at the beginning (0 h) and the end (8 h) of forskolin and DMSO treatment. Scale bar: 200 μm. Bottom: Quantification of forskolin-induced organoid swelling. TKA organoids were exposed to DMSO or forskolin for the indicated time periods. Normalized changes in organoid diameter were calculated by first computing at each time point and for each organoid under consideration the increase in diameter relative to the corresponding value at t=0 h, followed by normalization of forskolin-induced relative changes in diameter to those of DMSO-treated control samples. At least five organoids treated with DMSO or forskolin were analyzed per biological replicate and organoid line. **e**, Left: set-up for cultivating organoids in type I collagen at an air-liquid-interface and representative H&E stainings of organoid displaying different histological features (line 815). Scale bar: 200 μm. Right: Quantification of organoids following histological classification (morib.: moribund; dyspl.: dysplastic). Quantitative experiments in (**d** and **e**) were performed with three floxed/TKA organoid lines (815: n=3; 931: n=3; 947: n=3). Dots represent results of independent biological replicates and dot color identifies the organoid lines.

### TGFβ1 triggers collective invasion of TKA organoids and ECM remodeling

TGFβ1 is a prototypical tumor microenvironmental signal and prime inducer of EMT. Because the vast majority of human metastatic CRCs possess an intact TGFβ signaling machinery (Supplementary fig. 2a), we investigated the cellular and molecular consequences of treating organoids with TGFβ1. Whereas floxed organoids died when exposed to TGFβ1, TKA organoids were resistant to TGFβ1-induced cell death (Supplementary fig. 2b), probably due to expression of oncogenic Kras^G12D^ (30). Although levels of cleaved caspase-3, indicative of ongoing apoptosis, increased in TGFβ1-treated TKA organoids, they were even higher in solvent-treated organoids, and cleaved caspase-3 was detectable only in cells that had been shed into the lumen of solvent or TGFβ1-treated TKA organoids (Supplementary fig. 2c, d). Therefore, TGFβ1 does not appear to cause cell death in TKA organoids. Rather, TGFβ1 triggered massive morphological changes of TKA organoids which progressively lost their cystic shape, flattened, and extended multicellular protrusions, culminating in the formation of large, cohesive cell sheets and extended strands of cells which infiltrated the surrounding Matrigel (Fig. 2a; Supplementary movie 1). Boyden chamber assays independently confirmed acquired invasiveness of TGFβ1-stimulated TKA organoids (Fig. 2b, c). TKA organoids embedded in a type I collagen matrix and exposed to TGFβ1 also remained viable, changed their shape, and became invasive (Fig. 2d). Moreover, colon-derived TKA organoids responded to TGFβ1 treatment with highly similar morphological changes and also displayed a collective mode of invasion (Supplementary fig. 2e-g).

**Fig. 2:**
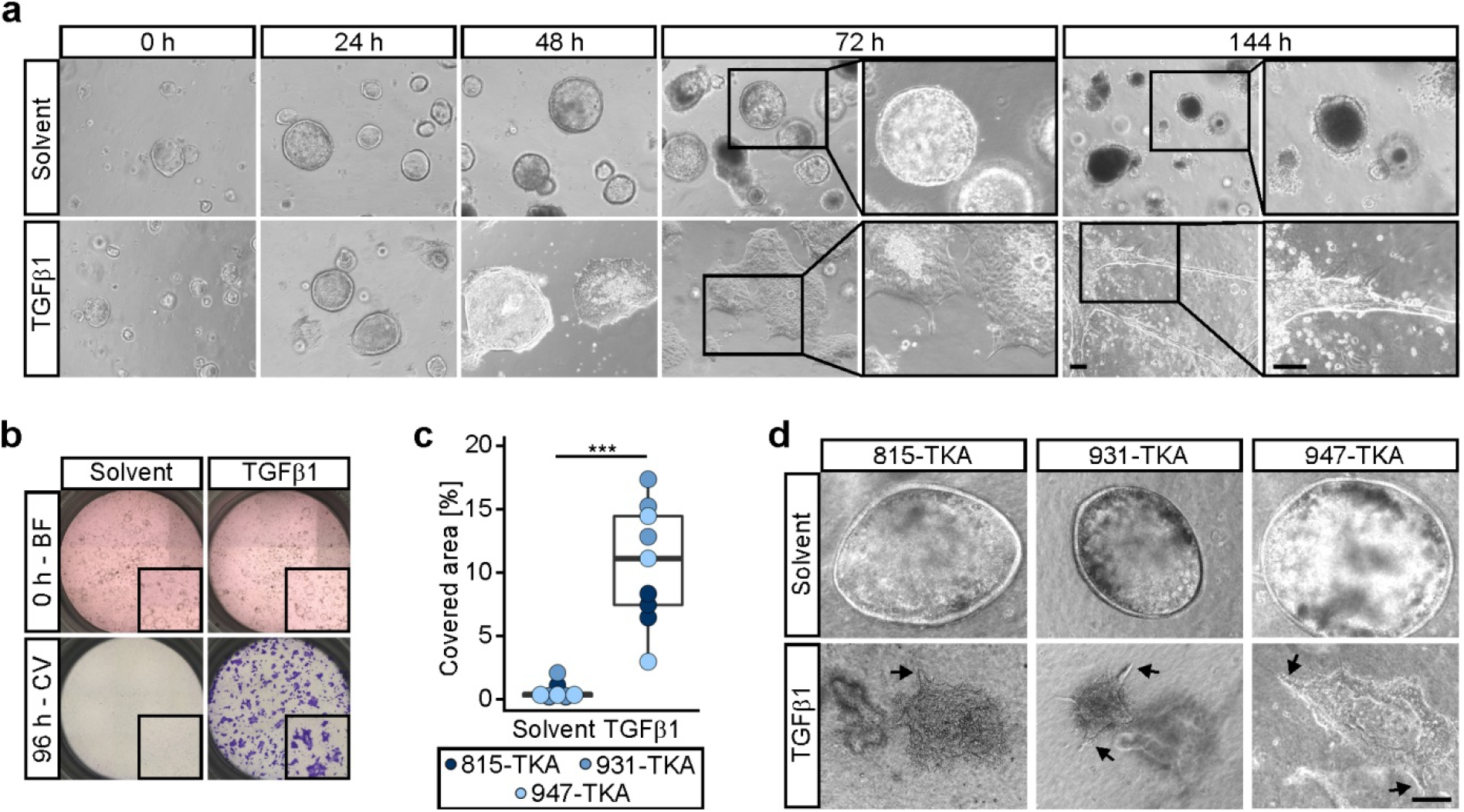
TGFβ1 triggers collective invasion of TKA organoids. **a**, Morphological appearance of TKA organoids (line 815) cultured in 3 mg/ml Matrigel and treated with solvent or TGFβ1 for the indicated time periods. Boxed areas are shown at higher magnification on the right. Similar TGFβ1-induced morphological changes were observed with TKA organoid lines from five different founder mice. Scale bars: 100 μm. **b**, Boyden chamber invasion assays with TKA organoids (line 931) seeded in 3 mg/ml Matrigel. Top: bright field (BF) images taken at 0 h of TGFβ1 and solvent treatment. Inserts show magnified views of the upper chambers. Bottom: crystal violet (CV) staining of invaded cells after 96 h of treatment. Inserts show magnified views of the bottom faces of the Boyden chambers. **c**, Quantification of invasion experiments as shown in (**b**) performed with TKA organoid lines 815 (n=3), 931 (n=3), and 947 (n=3). Each dot represents the result of a single invasion assay while dot color identifies the organoid lines. \*\*\**p=*0.0004; Mann-Whitney *U* test. **d**, Phase contrast microscopy of TKA organoids cultured in type I collagen in presence of solvent or TGFβ1. Arrows point at sites of collective invasion. Independent experiments were performed in three different organoid lines (815: n=3; 931: n=3; 947: n=3). Scale bar: 100 μm.

Dual fluorescent labelling with membrane-bound mTomato and nuclear H2B-GFP and confocal microscopy revealed that TGFβ1-treated TKA organoids adopted a dome-like structure with a central lumen and a broad circumferential rim of cells which coherently expanded outwards (Fig. 3a). Single cell delamination was not observed. Invading cell sheets formed only at the organoid base close to the surface of the cell culture plates. Cells at the invasive organoid perimeter produced spike-like fibers of fibronectin [involved in ECM assembly (31)], showed punctiform staining of the integrin β3 subunit [involved in adhesion to fibronectin (32)], and of vinculin [marking focal adhesions (33)] (Fig. 3b; Supplementary fig. 3). The observed topological restriction of invasion and formation of fibronectin spikes contrasts with the nuclear localization of Smad2/3 and, hence, active TGFβ signaling, and expression of fibronectin also in organoid cells constituting the dome structure (Supplementary fig. 4). This suggests that invasive behavior aside from TGFβ pathway activation additionally depends on a permissive microenvironment. Indeed, to allow uniform TGFβ1-induced phenotypic switching in organoid cultures we had to lower the Matrigel concentration. Conversely, organoid cells contracted the surrounding ECM while infiltrating: TGFβ1-treated TKA organoids grown in a type I collagen matrix promoted the formation of larger and more parallel aligned collagen bundles which could be visualized by picrosirius red staining (Fig. 3c). Altogether, TGFβ1 turned out to promote collective invasion of oncogenically transformed intestinal organoids which appears to involve reciprocal organoid/ECM interactions and ECM remodeling.

**Fig. 3:**
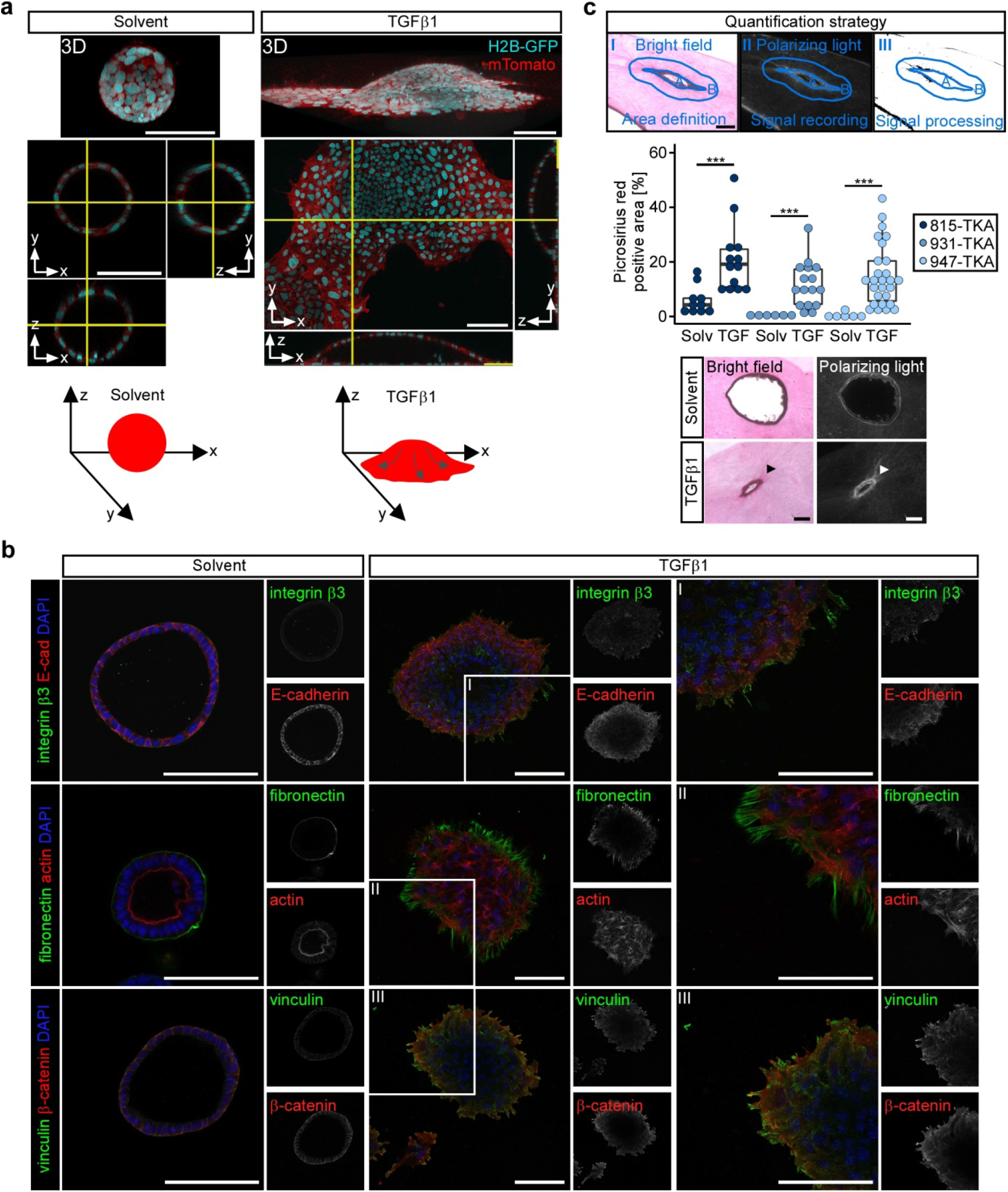
TGFβ1-stimulated TKA organoids interact with and remodel the ECM. **a**, TKA organoids (line 815) expressing nuclear H2B-GFP and membrane-bound mTomato were seeded in 3 mg/ml Matrigel, treated with solvent or TGFβ1 for 48 h, and subjected to live imaging using confocal microscopy. Displayed are 3D reconstructions and orthogonal views of organoid cross-sections (yellow lines: positions of the cross-sections along the x-, y- and z-axes). Scale bars: 100 μm (n=3). Bottom: schematic representations of TKA organoid morphologies. Arrows: proposed streaming of cells. **b**, Whole mount immunofluorescence staining and confocal microscopy of TKA organoids (line 815) seeded in 3 mg/ml Matrigel and treated with solvent or TGFβ1 for 72 h. Organoids were stained with antibodies against the indicated antigens. Actin was visualized by phalloidin staining. Nuclei were labeled using DAPI. Boxed areas are shown at higher magnification on the right. Pictures are representative for results obtained with three TKA organoid lines (815: n=1; 931: n=1; 947: n=1). Scale bars: 100 μm. **c**, Top: strategy for quantification of picrosirius red staining: (I) organoids were visualized by bright field illumination and the outline of the organoids (A) and a surrounding area (B) with an approximate width of 65 μm were marked. (II) Collagen was stained by picrosirius red and larger or parallel collagen bundles were visualized by polarizing light and signal intensities across the entire image were recorded. (III) For the final quantifications, only signals within area B and exceeding a defined threshold were considered. To allow comparison of organoids with different sizes, signal intensities were normalized to the size of area B. Middle: quantifications of picrosirius red-stained TKA organoids cultured in type I collagen and treated with solvent (solv) or TGFβ1 (TGF) for 96 h. Colored dots represent individual measurements from at least two independent experiments. Mann-Whitney *U* test. TKA organoid line 815: \*\*\**p=*0.0007 (solvent: n=10, TGFβ1: n=13); TKA organoid line 931: \*\*\**p=*0.0005 (solvent: n=6, TGFβ1: n=16); TKA organoid line 947: \*\*\**p=*0.0002 (solvent: n=6, TGFβ1: n=27). Scale bar: 100 μm. Bottom: exemplary pictures of TKA organoids (line 815) treated with solvent or TGFβ1.

### Canonical TGFβ signaling induces pEMT in oncogenically transformed organoids

Classically, TGFβ signaling induces complete EMT and single cell invasion (8, 10, 12, 27). To better understand TGFβ1-driven collective invasion of TKA organoids, we performed time-resolved gene expression analyses. TGFβ1 markedly induced expression of EMT TFs (Snail1, Zeb1), mesenchymal markers (fibronectin/*Fn1*, N-cadherin/*Cdh2*) (Fig. 4a, b), and *Itga5*, which, like *Itgb1*, is known to be upregulated during EMT (34). Significantly, integrin α5β1 is a fibronectin receptor (32) and promotes cancer cell migration and invasion (34–36). Western blot detection of phosphorylated Smad2/3 additionally confirmed TGFβ pathway activation. Surprisingly, TGFβ1 treatment did not diminish but rather increased expression of epithelial markers (E-cadherin/*Cdh1*, Ephb3, Foxa1), (Fig. 4a, b). Lastly, colonic TKA organoids responded to TGFβ1 treatment in a virtually identical fashion, showing concomitant upregulation of mesenchymal and epithelial gene expression and elevated levels of EMT-related integrins (Supplementary fig. 5a, b).

**Fig. 4:**
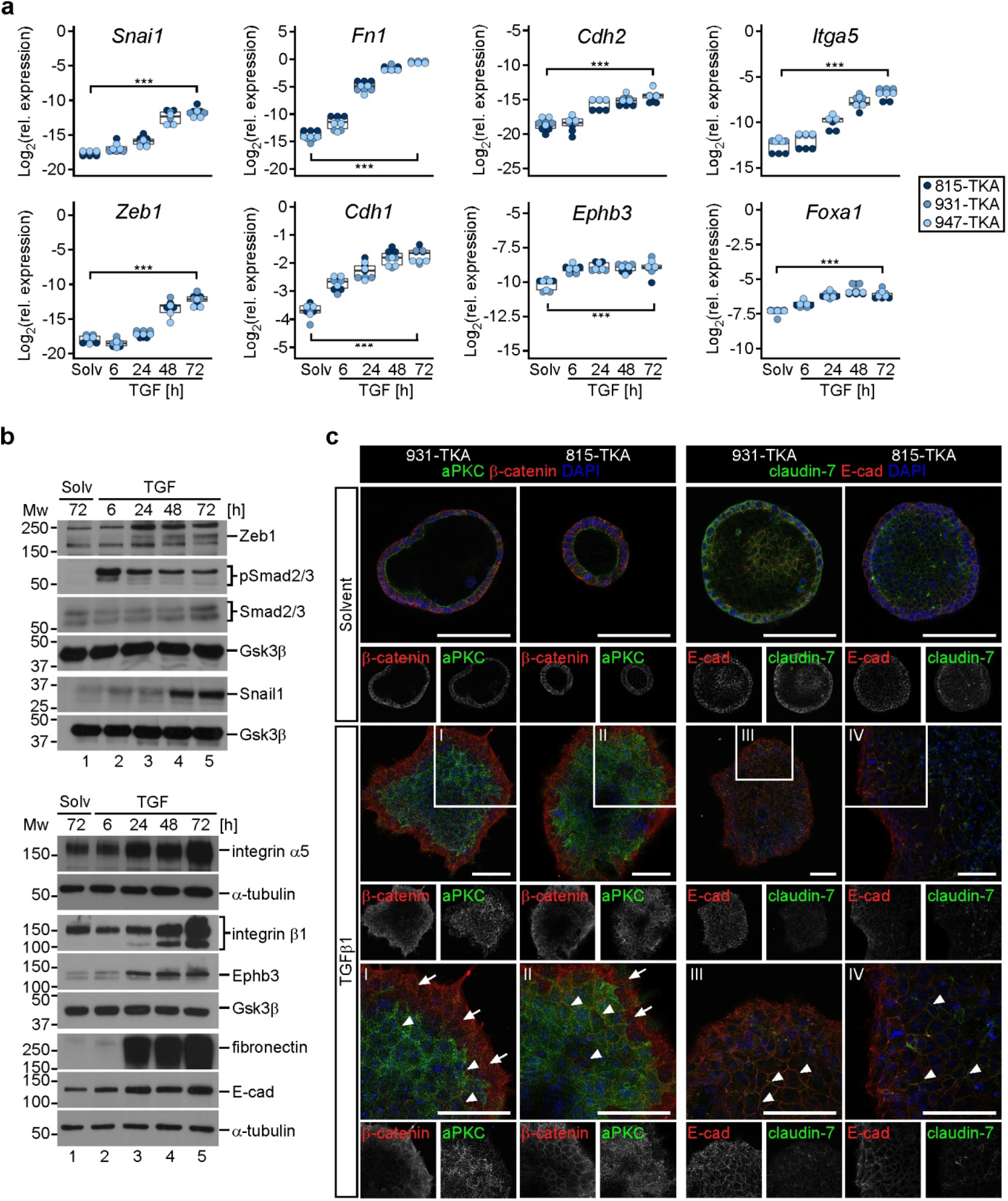
TGFβ1 induces a partial EMT in TKA organoids. **a**, Time-resolved gene expression analyses of TKA organoids seeded in 3 mg/ml Matrigel and treated with solvent (solv) or TGFβ1 (TGF) for the indicated time periods. RNA levels of EMT-TFs and EMT-associated genes were quantified by qRT-PCR and normalized to transcript levels of *Eef1a1*. Each dot represents the result of a single measurement while dot color identifies the organoid lines. Three independent biological replicates were performed for each organoid line (815: n=3; 931: n=3; 947: n=3). \*\*\**p*<0.001; statistical significance was analyzed using a linear regression model combined with Bonferroni correction for multiple comparisons. Exact *p*-values are provided in Supplementary table 1. **b**, Western blot analyses of phospho-Smad2/3 (pSmad2/3), total Smad2/3, EMT-TFs, and EMT-associated genes in TKA organoids (line 815) treated as in (**a**). Gsk3β and α-tubulin were included as loading control. Molecular weights of size standards are given in kDa. Results are representative for experiments performed with three TKA organoid lines (815: n=2; 931: n=2; 947: n=2); E-cad: E-cadherin. **c**, Whole mount immunofluorescence staining and confocal microscopy of TKA organoid lines 815 and 931 seeded in 3 mg/ml Matrigel and treated with solvent or TGFβ1 for 72 h. Organoids were stained for E-cadherin, claudin-7, β-catenin, and atypical protein kinase C (aPKC). Arrowheads indicate membranous E-cadherin, claudin-7, and β-catenin in the central part of the TGFβ1-treated organoid. Arrows highlight reduced staining of aPKC in the peripheral region of TGFβ1-stimulated organoids. Boxed areas are shown at higher magnification below. Reduced staining intensities in some central regions of organoids might be caused by technical issues related to whole mount staining and confocal microscopy of organoids. Similar results were obtained with three different organoid lines (815: n=1, 931: n=1, 947: n=1). Scale bars: 100 μm.

Aside from transcriptional repression, a change in intracellular localization of cell-cell adhesion molecules may also contribute to EMT (10). However, immunofluorescence stainings showed that the adherence junction proteins E-cadherin and β-catenin were retained at cell-cell borders throughout TGFβ1-treated TKA organoids (Fig. 4c, Supplementary fig. 5c). Only cells at the invasive front showed some increase in cytoplasmic E-cadherin and β-catenin staining, delocalization of the tight junction protein claudin-7, and reduction of aPKC (Fig. 4c). Despite these graded phenotypic changes, we conclude that TGFβ1-stimulated TKA organoids adopt a pEMT state distinguished by concurrent exhibition of epithelial and mesenchymal characteristics and largely maintained membranous E-cadherin and β-catenin.

The unexpected induction of pEMT by TGFβ1 prompted us to investigate the underlying signal transduction mechanisms. Overexpression of constitutively active TGF-β receptor type 1 (TGFBR1CA) (37) fully recapitulated TGFβ1-induced morphological changes and invasive behavior of TKA organoids (Supplementary fig. 6a-c). Conversely, expressing dominant negative TGF-β receptor type 2 (TGFBR2DN) (38), and pharmacologically inhibiting TGFBR1 blocked TGFβ1-induced phenotypic alterations (Fig. 5a-d, Supplementary fig. 6d). Finally, we knocked out *Smad4*, and generated quadruple mutant TKAS organoids (Fig. 5e-g). Unlike one might have expected (15, 17, 18, 21), *Smad4*-deficiency did not promote invasiveness *per se* but abrogated the TGFβ1 response (Fig. 5h-j, Supplementary fig. 6e). We conclude that the observed pEMT of oncogenically transformed intestinal organoids is conferred by receptor-mediated, canonical TGFβ signaling.

**Fig. 5:**
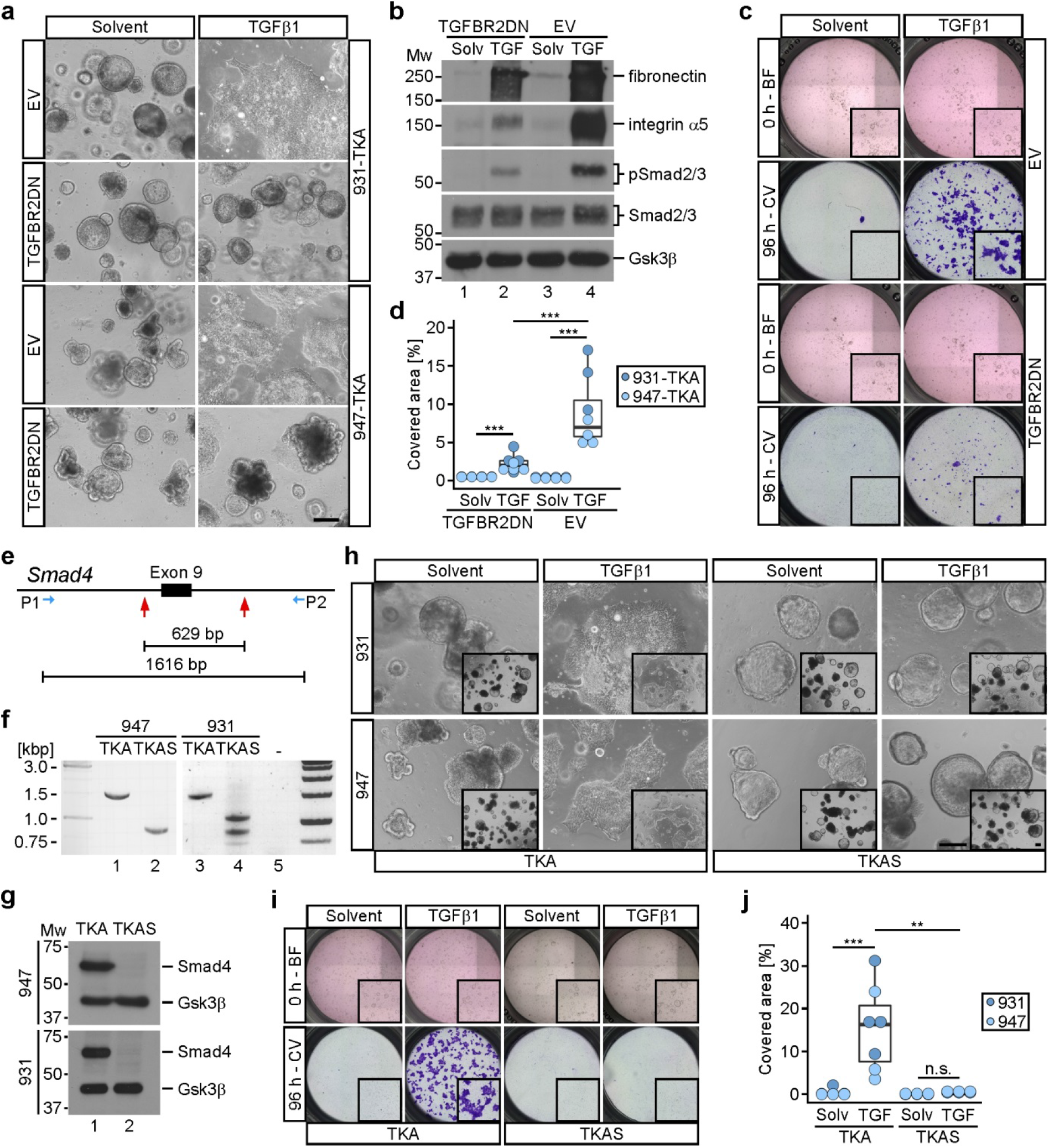
The TGFβ1 response in TKA organoids is mediated by canonical TGFβ-receptor/Smad signaling. **a**, Whole mount phase contrast microscopy of TKA organoids transduced with an empty vector (EV) or an expression vector for dominant negative TGFBR2 (TGFBR2DN), seeded in 3 mg/ml Matrigel, and treated with solvent or TGFβ1 for 72 h. Scale bar: 200 μm. **b**, Western blot expression analysis of the indicated proteins in control (EV) and TGFBR2DN-expressing TKA organoids (line 947) treated as in (**a**). Gsk3β detection served as loading control. Molecular weights of size standards are given in kDa. Panels (**a** and **b**) show representative results of two independent biological replicates for TKA organoid lines 931 (n=2) and 947 (n=2). **c**, Exemplary Boyden chamber invasion assays with control (EV) and TGFBR2DN-expressing TKA organoids (line 931) seeded in 3 mg/ml Matrigel. Bright field (BF) images were taken at 0 h of solvent and TGFβ1 treatment. Inserts: magnified views of the upper chambers. Invaded cells were visualized by crystal violet (CV) staining after 96 h of treatment. Inserts: magnified views of the bottom face of the Boyden chambers. **d**, Quantification of invasion experiments as shown in (**c**). Dots represent results of independent biological replicates (line 931: n=3; line 947: n=4). Dot color identifies the organoid lines. \*\*\**p*<0.001; Mann-Whitney *U* test. **e,** Scheme of the *Smad4* locus showing sgRNA target positions (red arrows) flanking exon 9 (black box) and the location of PCR primers used for genotyping. The distance between the sgRNA targets and the length of the PCR amplicon in *Smad4* wt organoids are given in base pairs (bp). **f**, Results of genotyping PCRs with genomic DNA from *Smad4* wt TKA and *Smad4* mutant TKAS organoid lines as indicated. Sizes of DNA standards are given in kilo base pairs (kbp). **g**, Western blot expression analyses for Smad4 in TKA and TKAS organoids. Gsk3β detection served as loading control (n=3). Molecular weights of size standards are given in kDa. **h**, Whole mount phase contrast microscopy of TKA and TKAS organoid lines seeded in 3 mg/ml Matrigel and treated with solvent or TGFβ1 for 72 h. Inserts display a larger field of view at lower magnification. Scale bars: 200 μm. **i**, Boyden chamber invasion assays performed with TKA and TKAS organoids (line 931) as in (**c**). **j**, Quantification of invasion experiments as shown in (**i**) performed with TKA and TKAS organoid lines 931 (n=4) and 947 (n=3). Dots represent results of individual experiments. Dot color identifies the organoid lines. \*\*\**p=*0.0006, \*\**p=*0.0012, n.s.: not significant (*p*=0.39); Mann-Whitney *U* test. For (**d** and **j**), exact *p*-values are provided in Supplementary table 1.

### Transcriptomes of TGFβ1-treated TKA organoids resemble CMS4 of human colorectal cancer

To comprehensively characterize TGFβ1-induced pEMT, we performed time-resolved transcriptome analysis by RNA sequencing (RNA-seq). Organoids were stimulated with TGFβ1 for up to 72 h or harvested at the onset of the experiment (C-0) and after 72 h of cultivation with solvent (C-72) to account for culture-dependent effects. Principal component (PC) analysis revealed high concordance between independent biological replicates from two organoid lines (Fig. 6a). A clear separation of control and TGFβ1-stimulated samples occurred along PC1, which accounts for most of the variances and likely reflects TGFβ1-induced gene expression changes over time. An additional segregation of samples along PC2 might be attributable to organoid maturation during cultivation (Fig. 6a). TGFβ1 caused extensive transcriptome changes eventually comprising 2,349 upregulated genes and 2,471 downregulated genes after 72 h of stimulation (Fig. 6b, Supplementary table 2; adjusted *p*-value <0.01, |log_2_(fold change[FC])|>1). Functional enrichment analyses of gene sets revealed gene ontology (GO) terms related to cell-cell adhesion, locomotion, extracellular structure organization, integrins, focal adhesion, and collagen formation as significantly enriched among genes upregulated by TGFβ1 (Fig. 6c). In agreement with this and with the observed changes in invasiveness, ECM deposition, and reorganization, the RNA-seq data showed TGFβ1-mediated elevated expression of ECM components, receptors, and remodeling enzymes (Supplementary table 3). Genes downregulated by TGFβ1 were enriched for GO terms connected to RNA and protein metabolic processes, DNA replication, and cell cycle (Supplementary fig. 7a). Indeed, compared to controls, TGFβ1-treated TKA organoids contained only few Ki67-positive cells which were concentrated in the organoid center (Supplementary fig. 7b). Accordingly, reduced proliferation may represent a common feature of partial and complete EMT (39).

**Fig. 6:**
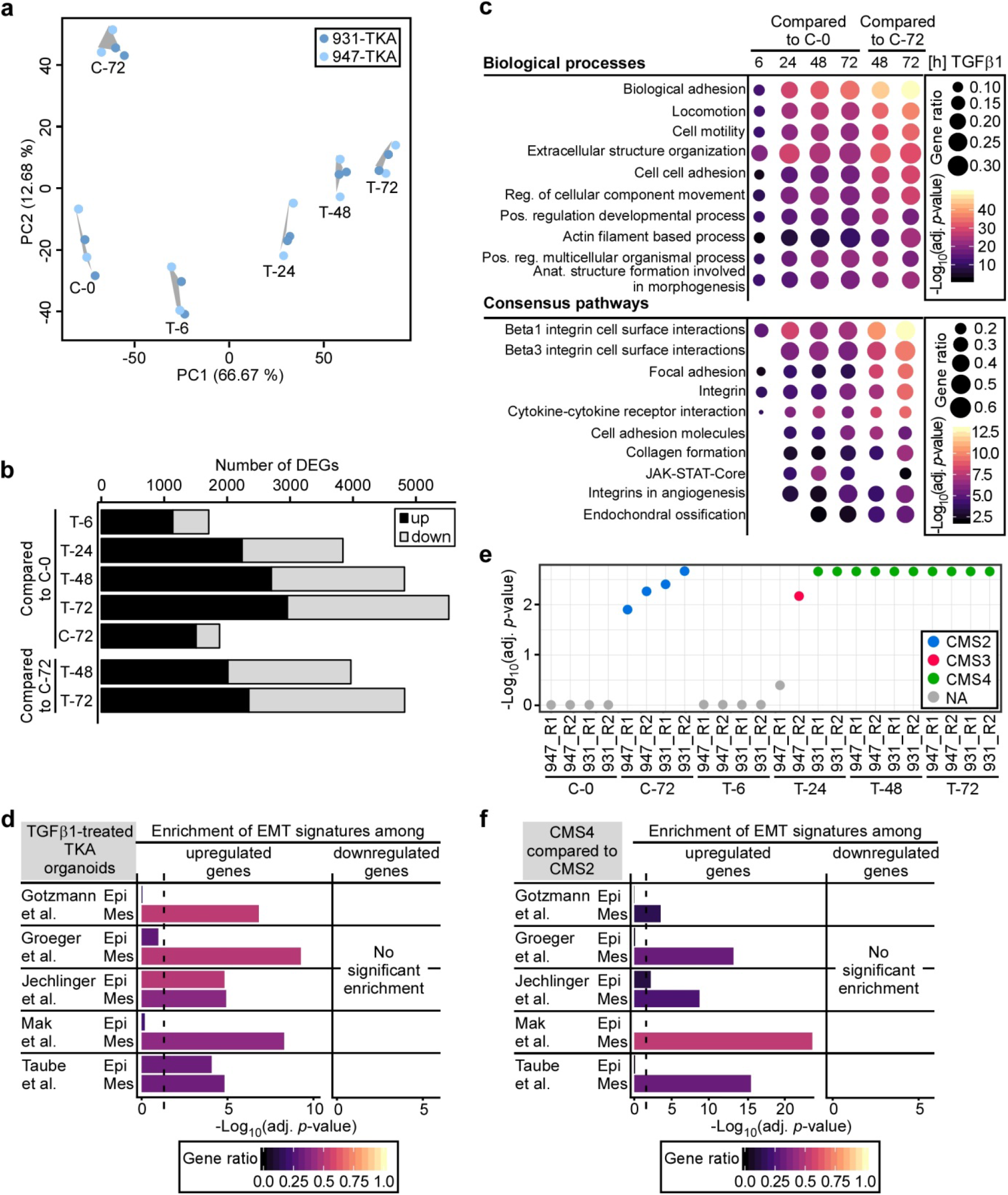
TGFβ1-induced global transcriptional deregulation features a partial EMT in TKA organoids. **a**, Principal component analysis (PCA) of RNA-seq data from TKA organoids seeded in 3 mg/ml Matrigel and treated with solvent (C) or TGFβ1 (T) as indicated. Results are based on two independent biological replicates for organoid lines 931 (n=2) and 947 (n=2). **b**, Numbers of differentially expressed genes (DEGs) were determined by performing pairwise comparisons of transcriptomes from TKA organoids harvested at the onset of the experiment (0 h of cultivation; C-0), cultivated for 72 h in solvent (C-72), and treated with TGFβ1 for the indicated periods of time. Black and grey segments of the bars: up- and downregulated genes, respectively. **c**, Functional enrichment analysis of genes upregulated upon TGFβ1 treatment (adjusted *p*-value <0.01, log_2_(FC)>1). The top ten GO terms from the indicated categories are listed. Dot size: ratio of upregulated genes compared to all genes within a set. Dot color: −log_10_(adjusted [adj.] *p*-value) of the enrichments. **d**, Exclusive enrichment of mesenchymal components of published EMT signatures among genes significantly upregulated in TGFβ1-treated TKA organoids. The analyses were conducted for DEGs from TKA organoids treated with TGFβ1 for 72 h compared to cultivation for 72 h in solvent. Published EMT signatures were split into subsets comprising epithelial (Epi) and mesenchymal genes (Mes) and processed separately. The color encodes the gene ratio. Length of the bars depicts the −log_10_ of the adj. *p*-values. Dotted line: adj. *p*-value =0.05. **e**, The independent biological replicates (R1, R2) of the transcriptomes as described in (**a** and **b**) were assessed for resemblance to the four consensus molecular subtypes (CMS) of CRC. Colored dots and their positions depict the CMS type and the adj. *p*-value, respectively. NA: no significant classification possible. **f**, Functional enrichment analyses of EMT sub-signatures as described in (**d**) among genes differentially expressed in colon cancers classified as CMS4 and CMS2. The color encodes the gene ratio. Length of the bars depicts the −log_10_ of the adj. *p*-values. Dotted line: adj. *p*-value =0.05.

Additionally, we related our transcriptome data to previously defined EMT gene expression signatures (40–44). Published data sets were split into epithelial and mesenchymal components, and enrichment of sub-signatures among the TGFβ1 up- and downregulated genes in TKA organoids was examined. Notably, TGFβ1-upregulated genes were significantly enriched for all mesenchymal and even two epithelial sub-signatures (Fig. 6d), while TGFβ1-downregulated genes showed no enrichment at all, confirming on a larger scale that TGFβ1-stimulated TKA organoids chiefly retain epithelial gene expression while concomitantly acquiring a mesenchymal transcriptional profile.

Human colorectal tumors can be classified into four subgroups CMS1-4 with distinctive transcriptomic features (25). To assess whether TKA organoids could be allocated to any of these, all RNA-seq data sets from control and TGFβ1-treated TKA organoids were compared to the CMS transcriptional profiles. Control organoids (C-0) and organoids treated with TGFβ1 for 6 h could not be classified (Fig. 6e), perhaps because mechanical disruption and reseeding of organoids had erased any typifying gene expression. The C-72 control group possessed gene expression properties of CMS2 which agrees well with hyperactive Wnt signaling in *Apc*-deficient TKA organoids (25). In contrast, past 24 h of TGFβ1 treatment, TKA organoids showed a uniform association with CMS4 which is defined as mesenchymal with signs of increased TGFβ signaling and EMT (25). Notably, when we interrogated human colon cancer transcriptome data from The Cancer Genome Atlas (TCGA) with respect to enrichment of epithelial and mesenchymal EMT sub-signatures, we found that genes upregulated in CMS4 compared to CMS2 were significantly enriched for all mesenchymal sub-signatures. Again, downregulated genes displayed no enrichment for any sub-signature (Fig. 6f). This further highlights the similarity between transcriptomes of TGFβ1-treated TKA organoids and human CMS4 tumors, and additionally hints that CMS4 tumors likewise exhibit a pEMT.

Next, we used ISMARA (Integrated System for Motif Activity Response Analysis) (45) to deduce from RNA-seq data potential regulatory DNA sequence motifs and TFs, whose activity might drive pEMT of TGFβ1-treated TKA organoids. Motifs identified by ISMARA were ranked according to their z-values as a measure for the contribution of each motif to the observed transcriptional pEMT program. Top entries for motifs with increased activity upon TGFβ1 treatment pointed to an involvement of Sox proteins, Sp1, Krüppel-like factors, Tead3/4, Snail1, Snail2, Zeb1, Fos and Jun family members, Smad4, and the Ets TF family (Supplementary table 4), all of which were implicated in EMT processes before (4, 7). DNA sequence motifs with reduced activity refer among others to the Nfy, E2f, and Myb families of cell cycle regulators which fits to the TGFβ1-induced decrease in proliferation in organoids.

We further compared ISMARA results to a collection of DNA sequence motifs which were previously implicated in shaping transcriptional landscapes at different stages of EMT (6). Activities of DNA sequence motifs common to all EMT states (Jun, Ets, Runx, Nfi, Tead3/4) were slightly higher under TGFβ1-treatment compared to controls and remained largely constant over time (Supplementary fig. 8a, b), except for the Tead3/4 motif which exhibited a large net difference in activity due to strongly decreased activity under control conditions. Possibly, this results from organoid maturation. In contrast to elevated and constant activities of pan-EMT state motifs, DNA sequence motifs which had previously been associated with distinct epithelial, mesenchymal and intermediate EMT states in squamous cell carcinoma and breast cancer (6), presented mostly with low z-values and only small activity changes, especially when taking into account changes in motif activity related to cultivation time (Supplementary fig. 8c). This difference in DNA sequence motif activity argues that alternative sets of TFs drive EMT processes in intestinal cancer versus other tumor entities. This notwithstanding, constant and increasing activities of motifs which were reported to be active specifically in epithelial cancer cells (Sox2, Trp63, Grhl1, Tfap2c) and early and late pEMT states (Sp1, Trp63, Tfap2c), further support that TGFβ1-induced pEMT of TKA organoids is distinguished by a reinforcement of epithelial features along with the additional gain of mesenchymal characteristics.

### TGFβ1-induced pEMT occurs independently from EMT master regulators

Snail, Zeb, and Twist TF family members are thought to fulfil key functions in EMT processes (4, 46). Since *Snai1* and *Zeb1* were the only EMT-TF genes upregulated in TGFβ1-treated TKA organoids (Supplementary table 2), we examined their roles in TGFβ1-inducible pEMT. Both genes were targeted applying a dual sgRNA-mediated deletion strategy, and multiple, clonally derived wildtype (TKA-Snai1^wt^, TKA-Zeb1^wt^) and knockout (TKA-Snai1^KO^, TKA-Zeb1^KO^) organoid lines were obtained (Fig. 7a). Importantly, Snail1 and Zeb1 deficiencies did not impair TGFβ pathway activity as demonstrated by unabated Smad2/3 phosphorylation (Fig. 7b). Despite complete absence of Snail1 and Zeb1, however, TGFβ1-regulated expression of EMT-associated genes, as well as TGFβ1-induced morphological conversion and invasiveness were unaffected (Fig. 7b; Supplementary fig. 9a, b). To complement these loss-of-function experiments, we generated TKA organoids with doxycycline (Dox)-inducible expression of Snail1 and ZEB1 (Supplementary fig. 10a, b). While ZEB1 overexpression had no discernible impact on organoid morphology and invasiveness (Supplementary fig. 10c-e), TKA organoids expressing Snail1 lost their cystic shape, occasionally infiltrated the surrounding Matrigel, and exhibited some invasiveness in Boyden chamber assays, albeit much less pronounced compared to TGFβ1 stimulation (Supplementary Fig. 10c-e). Neither ZEB1 nor Snail1 overexpression reproduced TGFβ1-mediated gene expression changes (Supplementary fig. 10f). All-in-all, overexpression of Snail1 and ZEB1 did not mimic the TGFβ1 response of TKA organoids. Thus, classical EMT-TFs are neither sufficient nor required for TGFβ1-mediated pEMT and collective invasion.

**Fig. 7:**
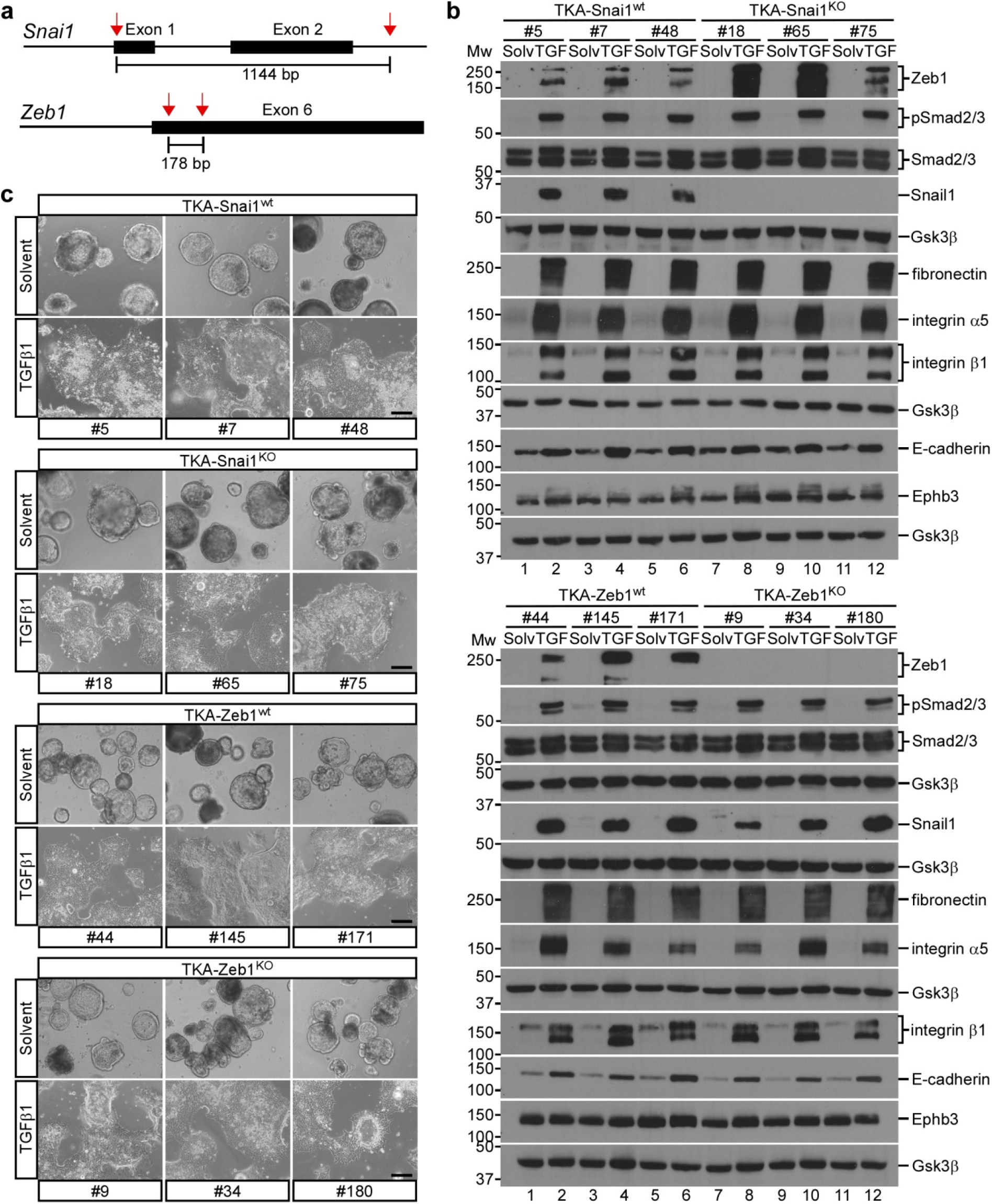
TGFβ1-induced collective invasion occurs independently from the EMT transcription factors Snail1 and Zeb1. **a**, Schematics of the *Snai1* and *Zeb1* loci showing sgRNA target positions (red arrows) and relevant exons (black boxes). The size of the expected deletions is given in base pairs (bp). **b**, Western blot expression analyses of the proteins indicated in TKA-Snai1^wt^, TKA-Snai1^KO^, TKA-Zeb1^wt^, and TKA-Zeb1^KO^ organoids derived from line 815 seeded in 3 mg/ml Matrigel and treated with solvent (solv) or TGFβ1 (TGF) for 72 h. Gsk3β detection served as loading control (n=3). Molecular weights of size standards are given in kDa. **c**, Morphological appearance of TKA-Snai1^wt^ (#5, #7, #48), TKA-Snai1^KO^ (#18, #65, #75), TKA-Zeb1^wt^ (#44, #145, #171), and TKA-Zeb1^KO^ (#9, #34, #180) organoids derived from line 815 cultured in Matrigel and treated with solvent or TGFβ1 for 72 h. Scale bars: 200 μm.

## Discussion

To decipher the molecular and cellular basics of tumor invasion and metastasis poses a persistent challenge, and it is not clear to which extent these processes are driven by genetic changes in cancer cells and by extrinsic factors from their surroundings. Here we used small intestinal and colonic organoids in a naïve, wildtype state to inflict oncogenic lesions only *in vitro*. This allowed us to assess the cell-autonomous impact of oncogenic transformation on epithelial integrity and invasiveness without prior exposure to any confounding influence from non-tumor tissue as e. g. in animal models. Under these circumstances, we found that the concomitant introduction of up to four mutations in *Apc*, *Kras*, *Trp53*, and *Smad4* was insufficient to elicit cell-intrinsic invasive behavior in organoids. This contrasts with results from genetically-engineered mouse models and organoid transplantation experiments where similar combinations of mutations led to the formation of invasive (since metastasizing) tumors (13–21). This difference in invasiveness *in vitro* and *in vivo* strongly argues that stimuli from the microenvironment were responsible for triggering tumor cell invasion in the *in vivo* settings. In this sense, oncogenic transformation constitutes an obligatory prerequisite, but only conditions tumor cells to become responsive to external pro-invasive signals. The implication of cell non-autonomous mechanisms in metastasis is consistent with the genetic similarity between primary and secondary lesions, and the failure to identify dedicated metastasis driver and suppressor genes in CRC (47). Furthermore, the critical importance of the tumor microenvironment for invasion and metastasis could readily explain inconsistent results concerning the mutational spectrum required for metastasis in autochthonous tumor models and upon heterotopic transplantations (13–21).

To test the idea that microenvironmental signals elicit invasiveness of oncogenically transformed organoids, we used the TGFβ pathway as a proof-of-principle. TGFβ signaling was previously shown to promote intestinal cancer metastasis, albeit indirectly by acting on non-cancer cells (22, 23). Although it is commonly pointed out that TGFβ pathway components are frequently mutated in colorectal tumors, and that disruption of TGFβ signaling promotes malignant progression in experimental models of intestinal cancer (15, 17–21) our results show that cancer cells themselves can be relevant targets of TGFβ signaling. In support of this, TGFβ receptors and SMAD genes are intact in more than 60% of human CRCs (24), and TGFβ pathway activity is evident in CMS4 cancers (25).

Our findings reinforce the importance of the tumor microenvironment for tumor invasion (48, 49). They further demonstrate that the tumor microenvironment can influence cancer cells in different, yet cooperating ways, first by providing an invasion-permissive ECM and second by supplying pro-invasive growth factors and cytokines. Nonetheless, oncogenically transformed organoids did not appear to just follow environmental cues, but actively participated in reciprocal interactions with their surroundings. The capacity to deposit and remodel ECM components in the tumor stroma therefore may not be restricted to cancer-associated fibroblasts and other non-cancer cells (50, 51).

TGFβ ligands are paradigmatic inducers of EMT through extensive transcriptional reprogramming which engages EMT-TFs and typically culminates in complete mesenchymal conversion (8, 10, 12, 27). Therefore, the ability of canonical, Smad4-dependent TGFβ signaling to induce pEMT without progression to a fully mesenchymal state represents a novel finding. Additionally, pEMT in TKA organoids was accompanied by collective invasion which likewise is not commonly associated with TGFβ-induced EMT (4, 5, 10, 27). Moreover, collective invasion and pEMT of TKA organoids were transcriptionally regulated whereby a mesenchymal gene expression program was implemented on top of a largely maintained epithelial program, including chiefly sustained membrane localization of E-cadherin and β-catenin. The hybrid epithelial/mesenchymal state of TGFβ1-treated TKA organoids therefore differs from previously described variants of pEMT which involved broadly downregulated epithelial gene expression or the abrogation of epithelial cell characteristics by transcription-independent internalization of cell adhesion molecules (6, 10). Interestingly, the gene regulatory network which was activated by TGFβ1 in TKA organoids, seemingly differs from that operating in pEMT states in squamous cell carcinoma and breast cancer (6, 11). Accordingly, we believe that the TGFβ1 response of oncogenically transformed intestinal cells represents a new and distinct pEMT program.

We suspect that the switch from TGFβ1-induced complete to partial EMT which we observed, is a consequence of context-dependent differences in the functionality of EMT-TFs. Not only were Snail1 and Zeb1 dispensable for TGFβ1-induced pEMT, but also overexpression experiments revealed much restricted EMT-inducing capacities of the two factors. Impaired function of EMT-TFs could be related to peculiarities of intestinal cells concerning posttranslational modifications of Snail1 and Zeb1 and to the expression of transcriptional co-factors (46). Above that, our results imply that EMT-TFs can be upregulated as passengers without functionally contributing to EMT processes. Conversely, lack of EMT-TF expression may not equal absence of EMT. This has implications for the use of Snail1 and Zeb1 as EMT markers, and for the interpretation of findings which refuted a role of EMT in metastasis based on knockout animals (52).

In agreement with their genetic constitution, TKA organoids exhibited a gene expression pattern resembling CMS2 which is dominated by Wnt pathway activity (25). Upon treatment with TGFβ1, organoids adopted a transcriptional profile highly similar to the transcriptome of CMS4 cancers which indeed are characterized by TGFβ pathway activation, mesenchymal features, high stromal content, and particularly poor prognosis (25). Convertibility to CMS4, thus, may not be limited to the sessile serrated adenoma path of colorectal carcinogenesis (26). Acquisition of CMS4 characteristics may only require a functional TGFβ pathway and could be determined primarily by microenvironmental influences. Notably, re-examination of gene expression patterns unearthed evidence for pEMT in human CMS4 samples, strengthening the significance of our findings for human cancer. Adding further to the pathophysiological relevance of our study, we note that collective invasion, which was triggered by TGFβ1 in TKA organoids, accurately reflects the predominant mode of stromal infiltration in most carcinomas.

Altogether, our study significantly expands the understanding of the mechanistic foundations of partial EMT and its context-dependent, distinctive appearances. Thereby, it may point out potential therapeutic strategies to eventually interfere with the metastatic cascade.

## Acknowledgements

The authors are grateful to the team of the Genomics and Proteomics Core Facility, German Cancer Research Center/DKFZ, Heidelberg, Germany, for next generation sequencing services, and to the staff of the Life Imaging Center (LIC) at the Center for Biological Systems Analysis (ZBSA) of the Albert-Ludwigs-University Freiburg for help with confocal microscopy. We especially appreciate the excellent support in image recording and analysis. We thank the members of the Hecht laboratory for helpful discussions and critical reading of the manuscript. We are particularly indebted to Vivien Freihen for her invaluable advice and initial introduction to the cultivation and handling of intestinal organoids. This study was supported in part by the Excellence Initiative of the German Research Foundation (GSC-4, Spemann Graduate School) and in part by the Ministry for Science, Research and Arts of the State of Baden-Wuerttemberg. Additional financial support was obtained from the Deutsche Forschungsgemeinschaft (DFG) (CRC-850 subprojects B5 to AH, B11 to AN, and Z1 to MB), and from the German Federal Ministry of Education and Research (BMBF) within the framework of the e:Med research and funding concept CoNfirm (FKZ 01ZX1708F to MB).

## Author Contributions

MF, AN, MB, and AH conceived and designed experiments. MF and MS generated data and analyzed all aspects of organoid morphology, histology, invasive behavior, and gene expression, and did the CRISPR/Cas9 genome editing experiments. MF and AN analyzed ECM remodeling by invasive organoids, acquired microscopic images and quantified collagen fiber formation. SD and MB carried out the bioinformatic analyses of the RNA-seq data, including identification of differentially expressed genes, gene set enrichment analyses, and CMS classification. MF, SD, MB, and AH interpreted data, created figures and wrote the manuscript. All authors critically read and commented on the final version of the manuscript.

## Materials and methods

### Organoid culture

Small intestinal and colonic organoid lines were established from C57BL/6N *Apc^580S/580S^*; *Kras^LSL-G12D/+^*; *Trp53^LSL-R172H/+^*; *tgVillin-CreER^T2^* mice (53–56) as described (57), and labeled with the identifier of founder animals. Mice were handled in accordance with legal regulations at the Center for Experimental Models and Transgenic Service of the University of Freiburg Medical Center (project registration number: X-17/07S). To recombine conditional alleles, organoids were treated with 0.5 μM 4-hydroxy-tamoxifen (4-OHT; Sigma Aldrich, H7904) for up to 120 h. Recombination was verified by PCR with primers listed in Supplementary table 5 and genomic DNA isolated with the ReliaPrep™ gDNA Tissue Miniprep System (Promega, A2052) and peqGOLD Tissue DNA Mini Kit (Peqlab, 12-3496-02). Upon inactivation of *Apc*, R-spondin-1 was omitted from the culture media.

### Manipulation of TGFβ and EGF signaling

For experiments involving TGFβ signaling, organoids were mechanically disrupted and seeded 30-40 h before treatments were started. To activate and inhibit TGFβ signaling, 5 ng/ml human TGFβ1 (Peprotech, 100-21) and 10 μM SB431542 (Selleckchem, S1067) were administered, respectively. To assess independence from EGFR signaling, organoids were mechanically disrupted and seeded in Matrigel two days before treatment with 0.8 μM Gefitinib (Selleckchem, S1025) for 72 h. Organoid viability was then determined by phase contrast microscopy and subsequent incubation with 500 μg/ml MTT (3-(4,5-dimethylthiazol-2-yl)-2,5-diphenyltetrazolium bromide) at 37°C for 1 h. Culture plates were imaged with a CanoScan 9950F scanner (Canon). Culture media supplemented with growth factors, inhibitors and other reagents were refreshed every 48 h.

### Assessment of epithelial integrity

To test for epithelial integrity, organoids were mechanically disrupted and seeded in Matrigel two days before treatment with 5 μM forskolin (Selleckchem, S2449). Swelling was monitored for 8 h at 20 min intervals using the JuLI™ Stage Real-Time Cell History Recorder (NanoEntek) and a 10x objective.

### Viral transduction

Lentiviral and retroviral particles were produced by co-transfecting HEK293T cells with viral vectors and packaging plasmids at a mass ratio of 1:0.75:0.3 using FuGENE®6 (Promega, E2691). Virus-containing medium was filtrated (0.45 μm) 48 h post-transfection and centrifuged at 8,000*g* and 4°C overnight. Viral pellets were resuspended in culture medium supplemented with 8 μg/ml Polybrene (Merck, TR-1003-G) and 10 μM Y-27632 (Selleckchem, S1049). For infection, organoids were dissociated into single cells by incubation with Accutase at 37°C for 10 min, mixed with the viral suspensions, plated on a Matrigel bed, and incubated at 37°C for 8-9 h. After washing with PBS, attached cells were overlaid with Matrigel and provided with culture medium containing Y-27632. Floxed organoids additionally received 1 μM valproic acid (Sigma Aldrich, P4543) and 1 μM CT99021 (Selleckchem, S2924). Selection with antibiotics was started three days (floxed organoids) and two days (TKA organoids) after transduction, using 5 ng/ml blasticidin (Invitrogen, R210-01), 2 μg/ml puromycin (Sigma Aldrich, P7255), 500 μg/ml geneticin (G418; Thermo Fisher Scientific, 10131027) (floxed organoids), and 700 μg/ml G418 (TKA organoids). For Dox-inducible gene expression, organoids were transduced with pMSCV-rtTA2-PGK-eGFP-F2A-NeoR, followed by a second infection with pRetroX-tight-Snail1-HA-PuroR, pRetroX-tight-ZEB1-HA-PuroR, or pRetroX-tight-MCS-PuroR (58, 59). Expression of Snail1 and ZEB1 was induced with 1 μg/ml Dox (Sigma Aldrich, D9891). For 4-OHT-inducible gene expression, TKA organoids were transduced with pMSCV-loxP-BlastR-loxP-TGFBR1(T204D)-F2A-eGFP (coding for TGFBR1CA), pMSCV-loxP-BlastR-loxP-TGFBR2(Δcyt)-F2A-eGFP (coding for TGFBR1DN), and pMSCV-loxP-BlastR-loxP-eGFP. Expression of TGFβ receptor mutants was induced by treating organoids with 0.5 μM 4-OHT. In case of TGFBR2DN, this was done three days prior to the start of invasion and gene expression experiments. For fluorescent labeling and live cell imaging, TKA organoids were transduced with pLenti-SV40-mTomato-P2A-H2B-GFP-PuroR. All constructs used in the study and their parental vectors are listed in Supplementary table 6.

### Genome editing

*Smad4*, *Snai1*, and *Zeb1* were inactivated by frame-shift-inducing exon deletions. For this, suitable exons were targeted by two single guide RNAs (sgRNAs; target sites listed in Supplementary table 5) selected using CCTop (60). Expression cassettes for sgRNAs were generated using vectors from the MuLE system (Supplementary table 6). For *Smad4* deletion, 400 ng of each sgRNA expression plasmid and 700 ng pCAG-Cas9-turbo-RFP vector were transfected into floxed organoids as described (17), except that single cell suspensions were generated with Accutase and transfection was done with Lipofectamine^®^ LTX (Invitrogen, 15338100). Three days after transfection, *Smad4*-deficient organoids were selected for by adding 100 ng/ml murine BMP-4 (Peprotech, 315-27) to the culture media while omitting Noggin. *Smad4*-deficient organoids were treated with 4-OHT to generate quadruple mutant TKAS organoids. To inactivate *Snai1*, floxed organoids were transduced with pLenti‐Cas9‐T2A‐BlastR and selected with blasticidin. Cas9-expressing organoids were transduced with pLenti-Dest-Snai1-sgRNA1+2-eGFP-F2A-NeoR-loxP and selected with G418 for one week. Thereafter, 0.5 μM 4-OHT was administered for 5 days to induce excision of loxP-flanked DNA sequences including the sgRNA-expressing provirus. Organoids were dissociated and sparsely seeded to facilitate clonal outgrowth. Single cell-derived organoids were manually picked, expanded, and screened by PCR with primers listed in Supplementary table 5. *Zeb1* was inactivated similarly, except that TKA organoids were simultaneously transduced with pLenti‐Cas9‐T2A‐BlastR and pLenti-Dest-Zeb1-sgRNA1+2-eGFP-F2A-NeoR-loxP, co-selected with blasticidin and G418, and exposed to 4-OHT for 3 days.

### Boyden chamber invasion assay

Organoids were mechanically disrupted and seeded into inserts for 24-well plates (Falcon, 353097) with Matrigel 30-40 h prior to treatments as described in the figure legends. Invaded cells on the bottom surface of the membranes were fixed with 4% paraformaldehyde at room temperature (RT) for 10 min and stained with 0.1% (w/v) crystal violet in water at RT for 10 min. For quantification, membranes were imaged using the BZ-9000 fluorescence microscope (Keyence) and the proportion of the membranes covered with invasive cells was determined using ImageJ.

### Air-liquid interface (ALI) culture

For ALI cultures (19), organoids were incubated in Cell Recovery Solution (Corning, 354253) on ice for 1 h, washed in PBS, mechanically disrupted, and seeded in type I collagen into 24-well transwell filter inserts. After gelation, culture medium was added to the outer well. Medium was exchanged every three days. After ten days, cultures were fixed in 4% paraformaldehyde at RT overnight, paraffin-embedded, sectioned into 5 μm slices, stained with hematoxylin and eosin, and imaged using the BZ-9000 fluorescence microscope (Keyence).

### Culture in type I collagen

Type I collagen (Collagen I HC, rat tail, Corning, 354249) was diluted with cold PBS to a concentration of 3 mg/ml. 1/20 (v/v) Medium 199 (Sigma Aldrich, M0650) was added, the pH was adjusted with NaOH, and the matrix was incubated on ice for 1 h. Organoids were prepared as described for ALI cultures, resuspended in the collagen matrix, and seeded on pre-warmed plates. After gelation, culture medium was added. Two days later, treatment with solvent or TGFβ1 was started.

### Picrosirius red staining

After 96 h of solvent or TGFβ1 treatment, organoid cultures in type I collagen were fixed in 10% formalin at RT overnight, paraffin embedded, and sectioned into 40 μm slices. Picrosirius red staining was performed as described (61). Sections were imaged using an Axioplan2 fluorescence microscope (Zeiss) equipped with a polarizer, an analyzer, and an Axiocam camera. All pictures taken with polarized or non-polarized light were acquired with the same exposure times. To quantify the formation of parallel and larger collagen bundles, images were acquired under polarizing light with a 540 nm filter. After acquisition, picrosirius red-derived signals above a defined threshold were measured within a 65 μm wide area immediately surrounding an organoid under investigation, while excluding the organoid itself.

### Immunofluorescence staining of paraffin sections and whole mounts

For sectioning, organoids were fixed in 4% paraformaldehyde overnight at 3-4 days after seeding, paraffin-embedded, and sectioned into 5 μm slices. Sections were stained as described (62). For whole mount imaging, organoids were seeded into chambered coverslips (Ibidi, 80826), treated with solvent or TGFβ1 for 72 h, and fixed *in situ* with 4% paraformaldehyde at 4°C for 45 min. After permeabilization with 0.5% (v/v) Triton X-100 in PBS at 4°C for 10 min and quenching with 0.1 M glycine in PBS at RT for 15 min, samples were incubated with blocking buffer (PBS with 10% FCS, 0.2% Triton X-100, 0.05% Tween20, 0.1% [w/v] BSA) at RT for 1 h. Primary antibodies were diluted in blocking buffer and incubated at 4°C overnight. After washing with blocking buffer, organoids were incubated with 0.3 μM DAPI and secondary antibodies diluted in blocking buffer. For actin staining, 5 u ml^−1^ phalloidin CF555 (Biotium, 00040) were added together with the secondary antibodies. After washing with blocking buffer, organoids were mounted with 0.1% (w/v) n-propyl gallate (Sigma Aldrich, P3130) in PBS. Primary and secondary antibodies are listed in Supplementary table 7.

### Fluorescence microscopy

Images of immunofluorescence stainings of paraffin sections were acquired using an Axio Observer.Z1 fluorescence microscope with an ApoTome2 equipment (Zeiss). Whole mount immunofluorescence stainings were imaged with a LSM 880 confocal microscope (Zeiss) with an Achroplan IR 40x/0.8 W objective and laser wavelengths of 405, 488, and 561 nm, unless stated differently. From image stacks (1.04 μm step size) orthogonal views of cross sections were generated using ZEN 2.3 and ImageJ. For live imaging, TKA organoids expressing mTomato and H2B-GFP were seeded into chambered culture plates (Ibidi, 80416). During confocal microscopy (settings as above), organoids were kept at 37°C and 5% CO_2_ in a Tokai Hit Incubator. Orthogonal views of cross sections and 3D reconstructions from live imaging data were generated using Huygens Professional for deconvolution and Imaris.

### RNA isolation, qRT-PCR, and RNA-seq

RNA was isolated and cDNA was synthesized with the peqGOLD MicroSpin Total RNA Kit (Peqlab/VWR Life Science, 12-6831) and the qScript™ Flex cDNA Kit (Quantabio, 95049), respectively. For qRT-PCR, the PerfeCTa® SYBR® Green SuperMix (Quantabio, 95049) was employed (primers listed in Supplementary table 8) with amounts of cDNA equivalent to 10 ng and 20 ng RNA when PCRs were conducted in a CFX384 and CFX96 Touch Real-Time PCR Detection System, respectively, (Bio-Rad Laboratories). Following normalization to *Gapdh* or *Eef1a1* transcripts, gene expression data were logarithmically transformed to yield normally distributed values, and are presented as log_2_(2^−ΔCT^). For global transcriptome analysis, RNA was collected from TGFβ1-treated organoids after 6, 24, 48, and 72 h, and from solvent-treated controls after 0 and 72 h of cultivation and paired-end sequenced on an Illumina HiSeq4000 at the Genome and Proteome Core Facility of the German Cancer Research Center, Heidelberg, Germany. FASTQ files were trimmed for sequencing adapters and low-quality reads with Trimmomatic (63). Reads were aligned to the Ensembl genome GRCM38 and reads per gene were quantified using STAR (64). For statistical analyses the R/Bioconductor package edgeR was used (65). We matched Ensembl IDs with EntrezIDs. If multiple Ensembl IDs matched to more than one Entrez ID, the one with the largest inter-quartile-range across samples was kept. Genes with less than one count per million in at least 3 samples were filtered out. Data were normalized with TMM (trimmed mean of M-values). Tagwise dispersion was calculated using edgeR. EdgeR was also used to compare data of the time series of TGFβ1 treatment to the 0 h and 72 h solvent controls. Differentially expressed genes (DEG) were defined by an adj. *p*-value <0.01 and a |log_2_(FC)|>1. Fisher’s exact test for Gene Ontology biological processes (GO:BP) (66), ConsensusPathDB (67) and a selected group of EMT signatures (41, 43, 40, 44, 42) was applied for functional enrichment analysis of DEGs. Terms were considered to be significantly regulated at an adj. *p*-value <0.05. All RNA-seq data were deposited in the Gene Expression Omnibus with the accession code GSE156553.

### CMS classification

The R/Bioconductor package CMScaller was used to determine the CMS of a given sample (68). For organoid transcriptome data, mouse genes were first mapped to their human orthologs using the getHomoGeneIDs function from the R/Bioconductor package GeneAnswers with the direct mapping method (69).

### Analysis of colon cancer data

Colon cancer RNA-seq data were downloaded from TCGA firehose (https://gdac.broadinstitute.org). Following CMS classification, only CMS2 and CMS4 samples were further processed. For these, we matched Ensembl IDs with EntrezIDs and, if multiple Ensembl IDs were matched to more than one Entrez ID, the one with the largest inter-quartile-range across samples was kept. Genes with less than one count per million in at least 3 samples were filtered out. TMM normalization, tagwise dispersion, and statistical analysis using edgeR were performed similarly to described above. For comparison of CMS4 to CMS2 samples, DEGs were defined by an adj. *p*-value <0.01 and |log_2_(FC)|>1 as calculated with edgeR. DEGs were subjected to functional enrichment analysis using Fisher’s exact test for the selected group of previously defined EMT signatures. A signature was considered to be significantly regulated if the adj. *p*-value <0.05.

### Protein expression analysis

Organoids were incubated in Cell Recovery Solution and nuclear extracts were prepared by resuspending organoids in nuclear extraction buffer A (10 mM HEPES/KOH pH 7.9, 0.1 mM EDTA, 10 mM KCl, 1x Complete® [Roche, 1697498], 1 mM DTT, 1x phosphatase inhibitor cocktails 2 and 3 [Sigma Aldrich, P5726/P0044]). After incubation on ice for 15 min, 0.5% (v/v) NP-40 was added, organoid suspensions were shortly vortexed, and nuclei were pelleted by centrifugation at 4°C and 16,100*g* for 2 min. The cytosolic supernatant was collected, and the nuclear pellet was washed once with nuclear extraction buffer A. Thereafter, nuclei were resuspended in 20 mM HEPES/KOH pH 7.9, 400 mM NaCl, 1 mM EDTA, 1x Complete®, 1 mM DTT, 1x phosphatase inhibitor cocktails 2 and 3, and incubated at 4°C for 30 min with constant shaking. Nuclear extracts were cleared by centrifugation at 4°C and 16,100*g* for 10 min. Protein concentrations were determined with the BioRad DC™ Protein Assay (BioRad, 500-0113). Depending on the protein yield in a given experiment, 25-40 μg of protein was separated by SDS-PAGE and transferred to nitrocellulose for protein detection as described (70). Antibodies are listed in Supplementary table 7.

### Query of publicly available cancer genome data and ISMARA

Genetic alterations in components of the TGFβ pathway were determined in CRC samples (47) using cBioPortal (71, 72). TF motif activities were predicted by the web-based tool ISMARA (45). Motifs were collected and sorted according to their significance as determined by their z-values. Motif activity profiles for TFs related to different EMT states were extracted according to ref. (7).

### Statistics and software

Data were analyzed and visualized using GraphPad Prism 5 (GraphPad Software) and RStudio (73). For time-resolved targeted gene expression studies, statistical analyses were done using a linear model allowing for simultaneous statistical testing for different time points. Normal distribution of the data was assessed with the car package (74). We applied the lm () function and defined the solvent control as intercept to estimate effect sizes and calculate *p*-values based on t-statistics. Analyses were completed by Bonferroni correction for multiple comparisons with the same data. In experiments in which two populations were compared, statistical significance was assessed using the two-tailed Mann–Whitney *U* test with a confidence interval of 95%. Box plots were generated with ggplot2 (75) and display the median with the lower and upper quartile. Whiskers show 1.5 times the interquartile range. Final figures were assembled using Canvas X 2017 (Canvas GFX, Inc.).

