## Supplementary Figures for "Canonical TGFβ signaling induces collective invasion in colorectal carcinogenesis through a Snail1- and Zeb1-independent partial EMT"

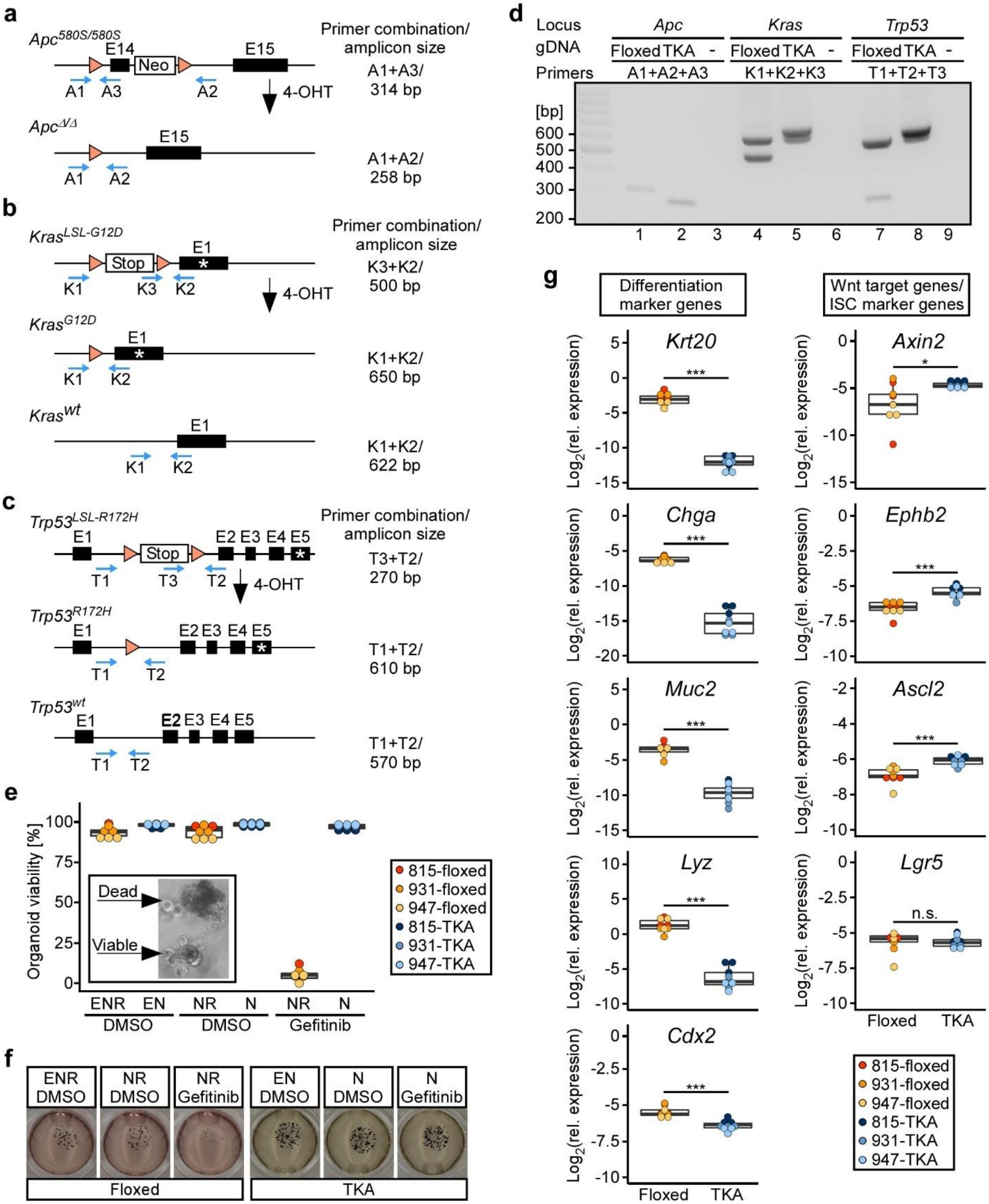


**Supplementary figure 1: *In vitro* characterization of oncogenically transformed organoids.** **a-c**, Schematic views of the *Apc*, *Kras*, and *Trp53* loci in small intestinal and colonic organoids derived from genetically engineered mice additionally carrying a *Villin-CreERT2* transgene. Black bars: exons (E) with locus-specific counting; orange triangles: LoxP elements; Neo:

Neomycin resistance gene; Stop: stop cassette preventing expression of floxed alleles; blue arrows: locus-specific PCR primers; asterisks: position of point mutations. Primer combinations used for genotyping and the expected amplicon lengths in base pairs (bp) for wildtype (wt), floxed, and recombined loci are listed to the right of the corresponding gene models. **a**, Genetically engineered mice carry two floxed alleles of the *Apc* gene. Treatment with 4-hydroxy-tamoxifen (4-OHT) leads to deletion of exon 14 and the adjacent Neo cassette ($Apc^{\Delta/\Delta}$). **b** and **c**, Genetically engineered mice are heterozygous at the *Kras* and *Trp53* loci with one allele being wildtype (wt) and the other being modified by insertion of floxed stop cassettes (LSL). Treatment with 4-OHT removes the LSL cassettes and results in expression of $Kras^{G12D}$ and $p53^{R172H}$. **d**, Gel electrophoresis separating DNA fragments generated by multiplex-PCRs with locus-specific primer combinations as indicated and genomic DNA (gDNA) isolated from floxed organoids and TKA organoids with recombined $Apc^{\Delta/\Delta}$, $Kras^{G12D}$ and $Trp53^{R172H}$ loci. Negative control samples received $H_20$ (-) as replacement for gDNA. **e**, Dependence of floxed and TKA organoids on EGFR signaling was determined by adding Gefitinib to the organoid culture media supplemented with the indicated combinations of EGF (E), Noggin (N), and R-spondin-1 (R). Treatment with DMSO served as solvent control. After 72 h, organoid viability was judged by microscopy. Examples for viable and dead organoids are shown in the box. **f**, Representative images of MTT staining of floxed and TKA organoids (line 815) treated as described in (**e**). **g**, Gene expression analyses of marker genes for differentiated intestinal cells, intestinal stem cells (ISC), and Wnt target genes by qRT-PCR. Expression was normalized to *Gapdh*. ***$p$<0.001, *$p$<0.05; Mann-Whitney *U* test. Exact *p*-values are provided in Supplementary table 1. For (**e-g**) three independent biological replicates were performed for each of three different floxed and TKA organoid lines (815: n=3, 931: n=3, 947: n=3) seeded in 7 mg/ml Matrigel. Dots represent results of individual experiments while dot color identifies the organoid lines.

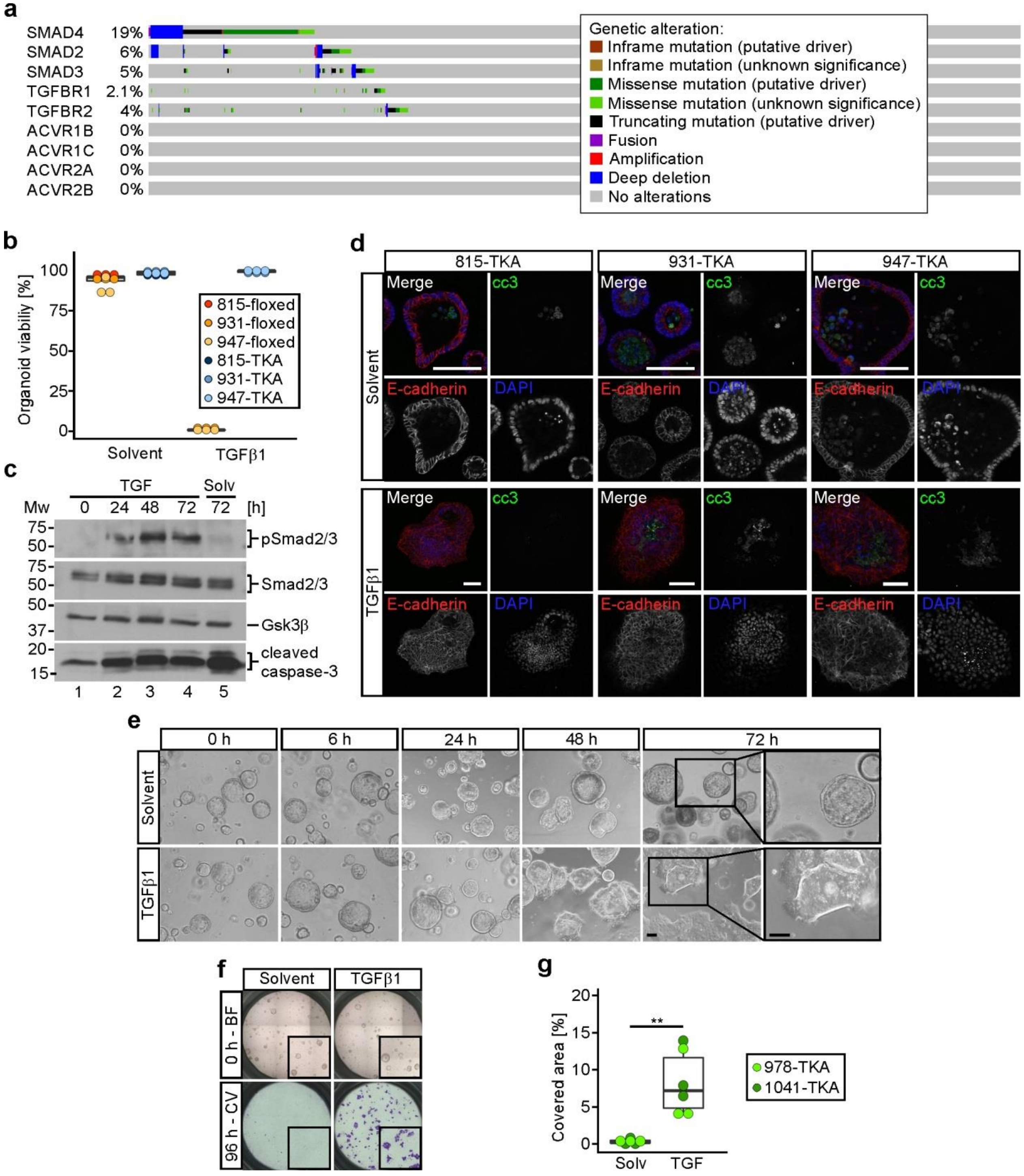


**Supplementary figure 2: Apoptosis resistance of oncogenically transformed small intestinal organoids and TGFβ1-induced collective invasion of colonic TKA organoids. a**, Nature and frequency of genetic alterations in components of the TGFβ signaling pathway in 1,134 CRC samples analyzed by cBioPortal. The majority (63.9%) of the examined CRC samples have no genetic alterations in the TGFβ signaling pathway. **b**, Floxed and TKA small intestinal organoids were seeded in 7 mg/ml Matrigel and treated with solvent or TGFβ1 for 72 h and organoid viability was judged by microscopy. Dots represent results of individual experiments while dot color identifies the organoid lines. Independent experiments were performed with three different floxed/TKA organoid lines, 815: n=3, 931: n=3, 947: n=3. **c**, Western blot analysis of cleaved caspase-3 in small intestinal TKA organoids (line 815) seeded in 3 mg/ml Matrigel and treated with solvent or TGFβ1 for the indicated periods of time. Phosphorylation of Smad2/3 was analyzed as indication for active TGFβ signaling. Gsk3β detection served as loading control. Molecular weights of size standards are given in kDa. **d**, Whole mount immunofluorescence staining of small intestinal TKA organoids seeded in 3 mg/ml Matrigel and treated with solvent or TGFβ1 for 72 h. Organoids were stained for cleaved caspase-3 (cc3) and E-cadherin. Nuclei were stained by DAPI. For (**c**) and (**d**), three independent biological replicates were performed with three different organoid lines (815: n=1; 931: n=1; 947: n=1). Scale bars: 100 µm. **e**, Morphology of colonic TKA organoids (line 978) cultured in 3 mg/ml Matrigel and treated with solvent or TGFβ1 for the indicated periods of time. Boxed areas are shown at higher magnification on the right. Scale bars: 100 µm. **f**, Boyden chamber invasion assays with colonic TKA organoids (line 1041) seeded in 3 mg/ml Matrigel. Top: bright field (BF) images taken at 0 h of solvent and TGFβ1 treatment. Inserts show magnified views of the upper chambers. Bottom: crystal violet (CV) staining of invaded cells after 96 h of treatment. Inserts show magnified views of the bottom faces of the Boyden chambers. **g**, Quantification of invasion experiments as shown in (**f**) performed with colonic TKA organoid lines 978 (n=3) and 1041 (n=3). Each dot represents the result of a single invasion assay while dot color identifies the organoid lines. Solv: solvent; TGF: TGFβ1. **$p$=0.0022; Mann-Whitney $U$ test.

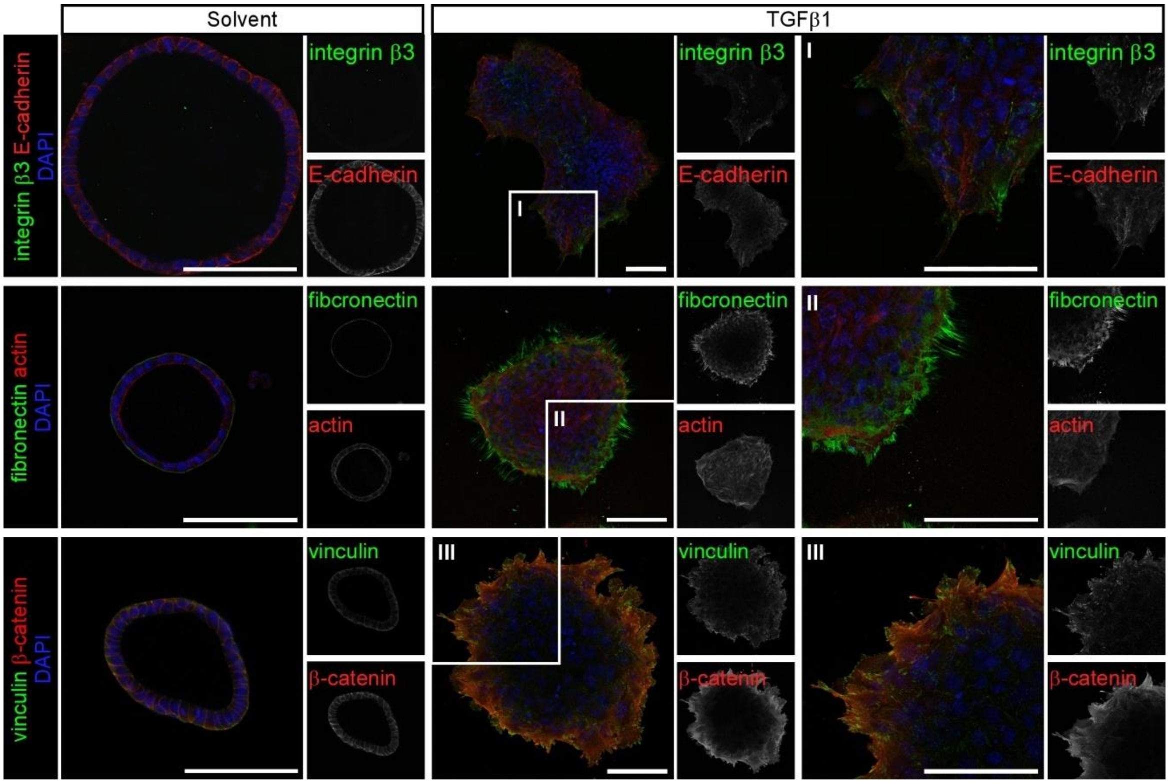


**Supplementary figure 3: Compartmentalization of TGFβ1-stimulated TKA organoids.** Whole mount immunofluorescence staining and confocal microscopy of TKA organoids (line 931) cultured in 3 mg/ml Matrigel and treated with solvent or TGFβ1 for 72 h. Organoids were stained with the indicated antibody combinations against integrin β3, E-cadherin, fibronectin, vinculin, and β-catenin. Actin was visualized by phalloidin staining. Nuclei were labeled with DAPI. Boxed areas I, II, and III are shown at higher magnification on the right. Pictures are representative for results obtained with three different TKA organoid lines (815: n=1, 931: n=1, 947: n=1). Scale bars: 100 µm.

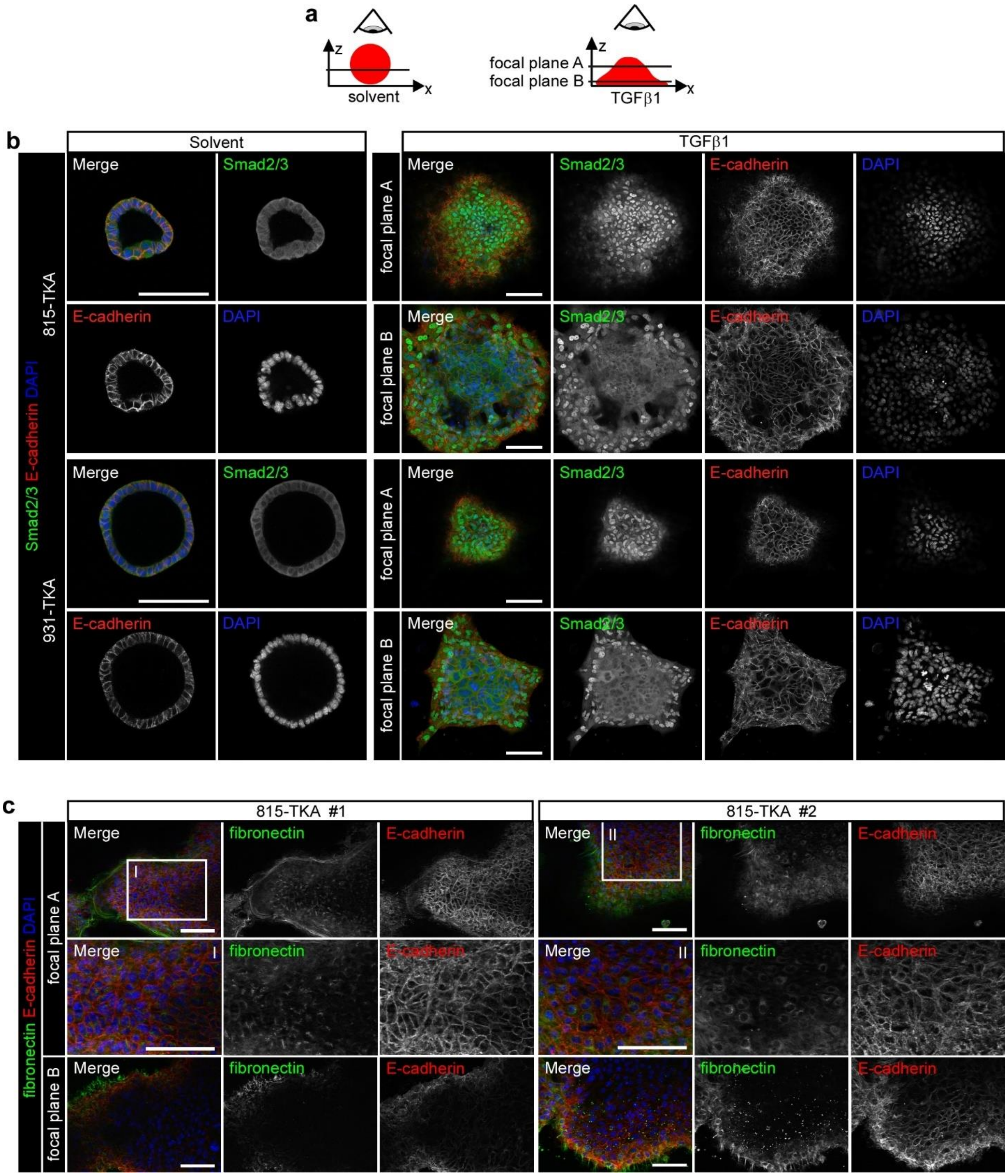


**Supplementary figure 4: Activation of TGFβ signaling and mesenchymal marker gene expression in the dome structure of TGFβ1-stimulated TKA organoids. a**, Direction of view and position of focal planes for images shown in (**b**) and (**c**). **b**, Whole mount immunofluorescence staining and confocal microscopy of TKA organoid lines 815 and 931 seeded in 3 mg/ml Matrigel and treated with solvent or TGFβ1 for 72 h. Organoids were stained for Smad2/3 and E-cadherin. Nuclei were visualized with DAPI. TGFβ1-stimulated organoids were imaged at focal planes within the dome structure (A) and at their base close to the bottom of the cell culture plates (B). **c**, Whole mount immunofluorescence staining and fluorescence microscopy of TKA organoids (line 815) treated as in (**b**). Organoids were stained for fibronectin and E-cadherin. Nuclei were labeled with DAPI. Boxed areas I and II in images of focal plane A are displayed at higher magnification in the panels underneath. Shown are two representative examples (line 815-TKA organoids #1 and #2) of three independent biological replicates, n=3. Images were acquired using an Axio Observer.Z1 fluorescence microscope with an ApoTome2 equipment (Zeiss). Scale bars: 100 µm.

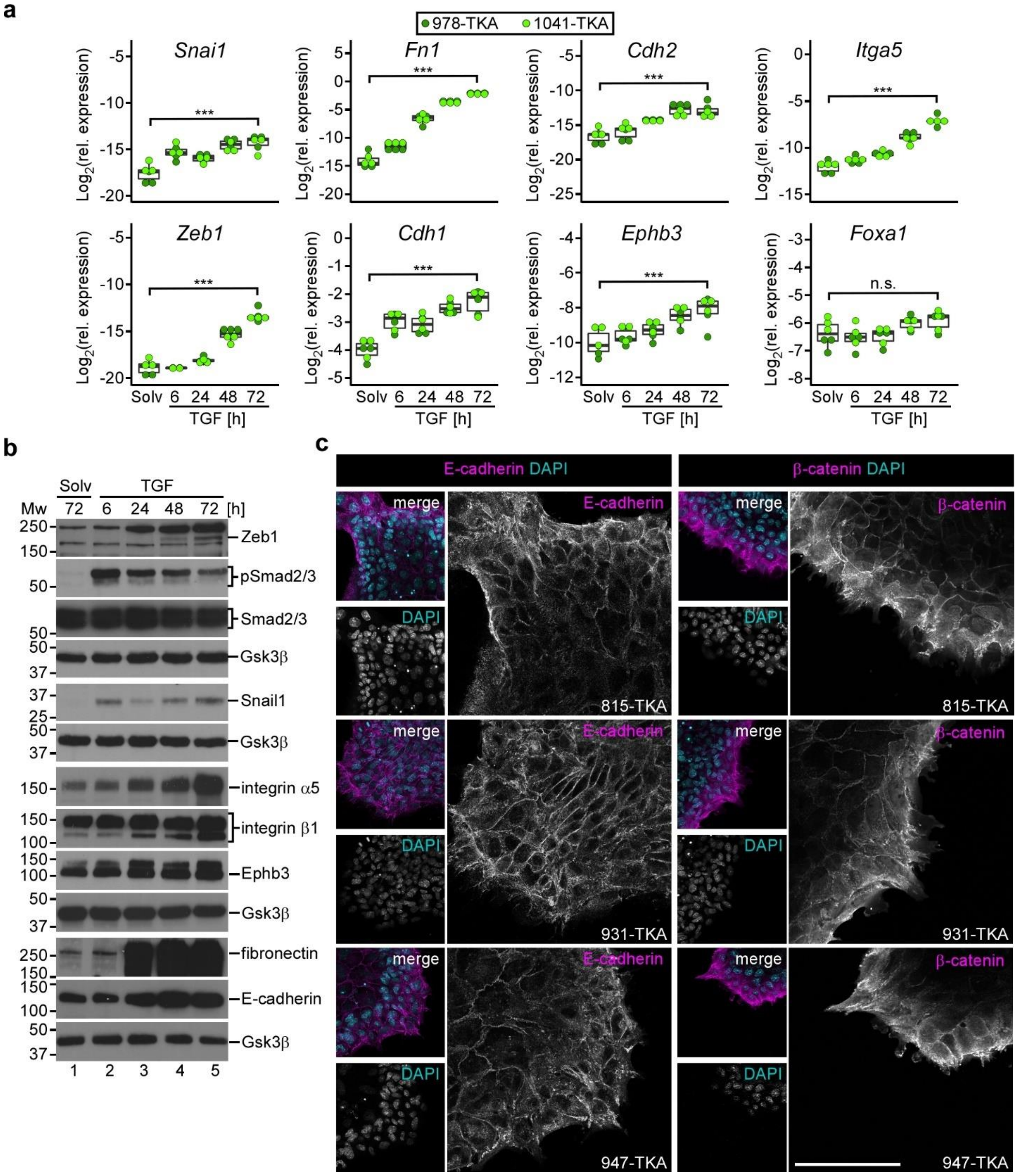


**Supplementary figure 5: TGFβ1 induces a partial EMT in oncogenically transformed small intestinal and colonic organoids. a**, Gene expression analysis of colonic TKA organoids seeded in 3 mg/ml Matrigel and treated with solvent (solv) or TGFβ1 (TGF) for the indicated periods of time. Gene-specific transcripts of EMT-TFs and EMT-associated genes were quantified by qRT-PCR and normalized to transcript levels of *Eef1a1*. Each dot represents the result of a single measurement while dot color identifies the organoid lines. Three independent biological replicates were performed for two organoid lines (978: n=3; 1041: n=3). ***$p<0.001$, n.s.: not significant; statistical significance was analyzed using a linear model combined with Bonferroni correction for multiple comparisons. Exact *p*-values are provided in Supplementary table 1. **b**, Western blot analyses of phosphorylated Smad2/3 (pSmad2/3), total Smad2/3, EMT-TFs, and EMT-associated genes in colonic TKA organoids (line 978) treated as in (**a**). Molecular weights of size standards are given in kDa. Gsk3β detection served as loading control (n=3). **c**, Additional examples for whole mount immunofluorescence stainings and confocal microscopy of TKA organoid lines 815, 931, and 947 treated with TGFβ1 for 72 h. Organoids were stained for E-cadherin or β-catenin. Nuclei were labelled using DAPI. Scale bar: 100 µm.

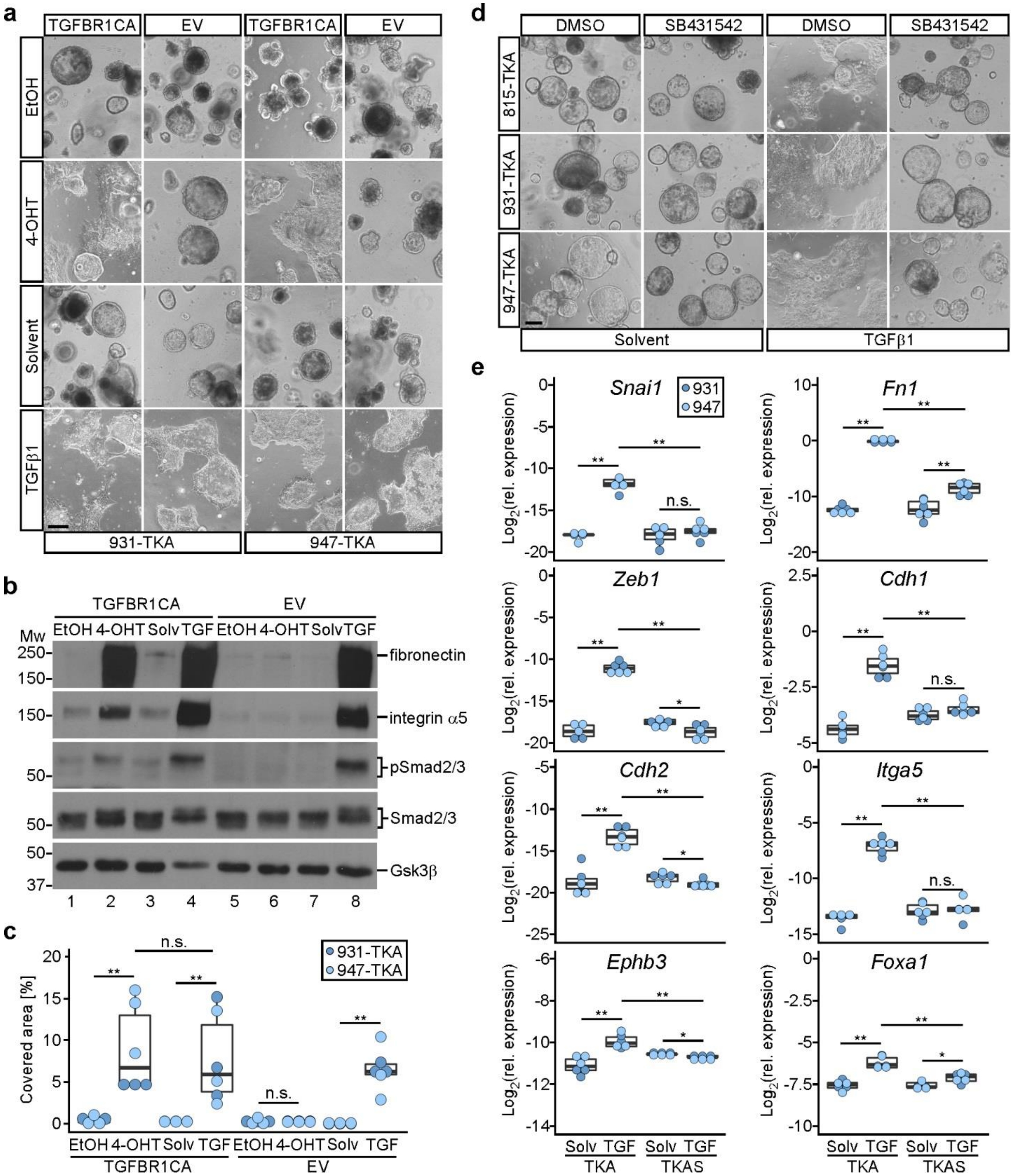


**Supplementary figure 6: The TGFβ1 response in TKA organoids depends on canonical TGFβ-receptor/Smad signaling. a-c**, TKA organoids, transduced with an empty vector (EV) or a vector for inducible expression of a constitutively active form of TGFBR1 (TGFBR1CA), were seeded in 3 mg/ml Matrigel and treated with ethanol (EtOH) or 4-OHT for 96 h to trigger TGFBR1CA production. Alternatively, organoids received solvent (solv) or TGFβ1 (TGF) for 72 h. **a**, Whole mount phase contrast microscopy of transduced TKA organoids. Two independent biological replicates were performed with two different organoid lines (931: n=2, 947: n=2) Scale bar: 200 µm. **b**, Western blot analyses of phosphorylated Smad2/3 (pSmad2/3), total Smad2/3, fibronectin, and integrin α5 in transduced TKA organoid line 947 treated as in (**a**). Gsk3β detection served as loading control. Molecular weights of size standards are given in kDa. Results shown are representative for three independent biological replicates performed with organoid lines 931 (n=3) and 947 (n=3). **c**, Quantification of Boyden chamber invasion assays performed with transduced organoids as described in (**a**). Three independent biological replicates were performed with each of two different TKA organoid lines (931: n=3, 947: n=3). Dots represent results of individual experiments. Dot color identifies the organoid lines. $^{**}p<0.01$, n.s.: not significant; Mann-Whitney *U* test. **d**, Whole mount phase contrast microscopy of TKA organoids seeded in 3 mg/ml Matrigel and treated with solvent or TGFβ1 in presence of DMSO or SB431542 for 72 h. Images shown are representative for three independent biological replicates performed with three different organoid lines (815: n=3, 931: n=3, 947: n=3). Scale bar: 200 µm. **e**, Gene expression analyses of *Snai1, Zeb1, Fn1*, *Cdh2*, *Itga5*, *Cdh1*, *Ephb3,* and *Foxa1* in TKA and TKAS organoids seeded in 3 mg/ml Matrigel and stimulated with solvent or TGFβ1 for 72 h. Gene-specific transcripts were quantified by qRT-PCR and normalized to transcript levels of *Eef1a1*. Three independent biological replicates were performed with TKA and TKAS organoids derived from lines 931 (n=3) and 947 (n=3). Dots represent results of individual experiments while dot color identifies the organoid lines. $^{**}p<0.01$, $^{*}p<0.05$, n.s.: not significant; Mann-Whitney *U* test. For (**c** and **e**), exact *p*-values are provided in Supplementary table 1.

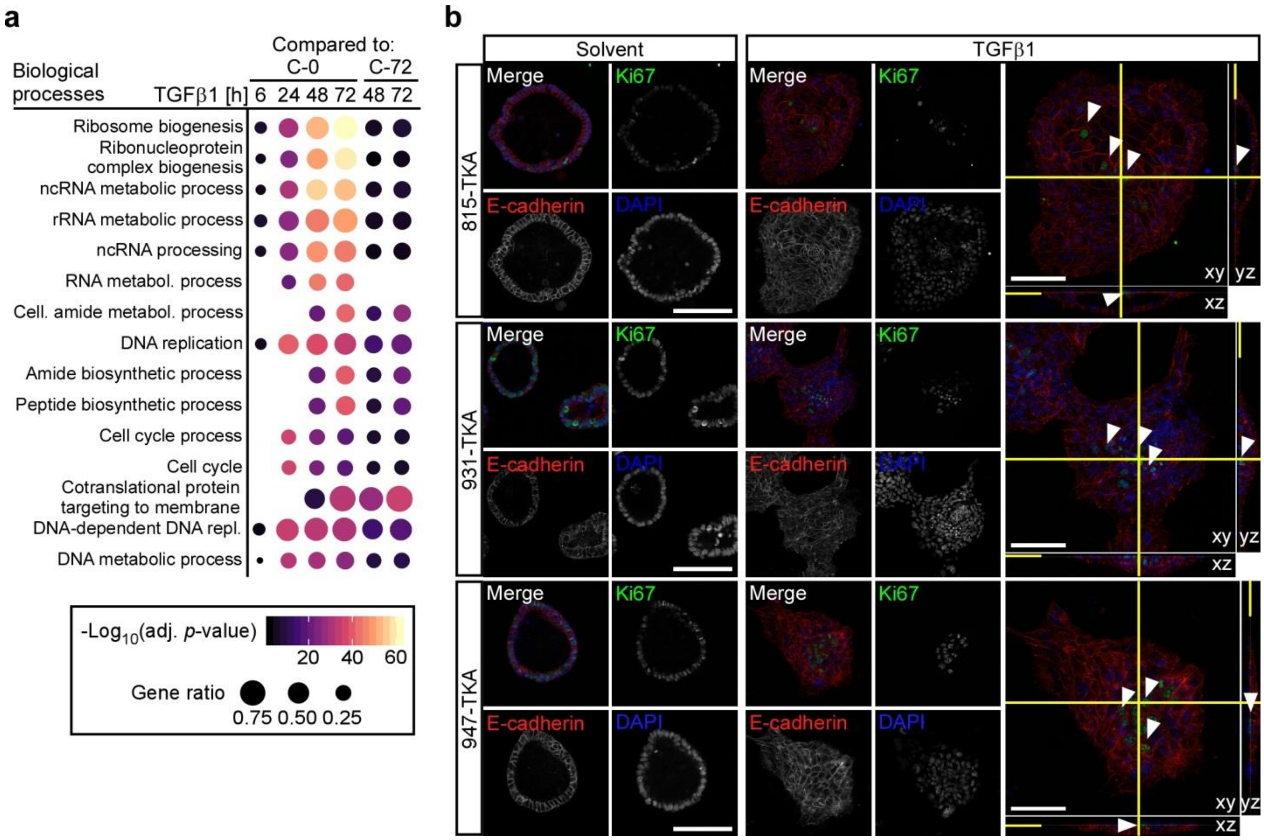


**Supplementary figure 7: TGFβ1-induced gene expression changes in TKA organoids indicate repression of proliferation.** **a**, Functional enrichment analysis of genes downregulated upon TGFβ1 treatment. Downregulated genes were determined by comparing transcriptomes of organoids treated with TGFβ1 for the indicated periods of time to those of organoids harvested immediately at the onset of the experiment (0 h of cultivation; C-0) or cultivated for 72 h in solvent (C-72) using an adjusted (adj.) *p*-value<0.01 and $\log_2$(FC)<-1 as thresholds. The top fifteen GO terms from the category "Biological processes" significantly enriched among downregulated genes are listed. Dot size reflects the ratio of downregulated genes compared to all genes in each GO term, while the color encodes the -$\log_{10}$(adj. *p*-value) of the enrichment. **b**, Whole mount immunofluorescence staining and confocal microscopy of TKA organoids seeded in 3 mg/ml Matrigel and treated with solvent or TGFβ1 for 72 h. Organoids were stained for Ki67 and E-cadherin. Nuclei were visualized with DAPI. Arrowheads mark a few remaining Ki67-positive cells in the center of TGFβ1-stimulated organoids. The right panel displays orthogonal views of cross-sections of TGFβ1-stimulated organoids. Yellow lines indicate the positions of the cross-sections along the x-, y-, and z-axes. Images show representative results of two independent biological replicates obtained with the different organoid lines indicated (n=2). Scale bars: 100 µm.

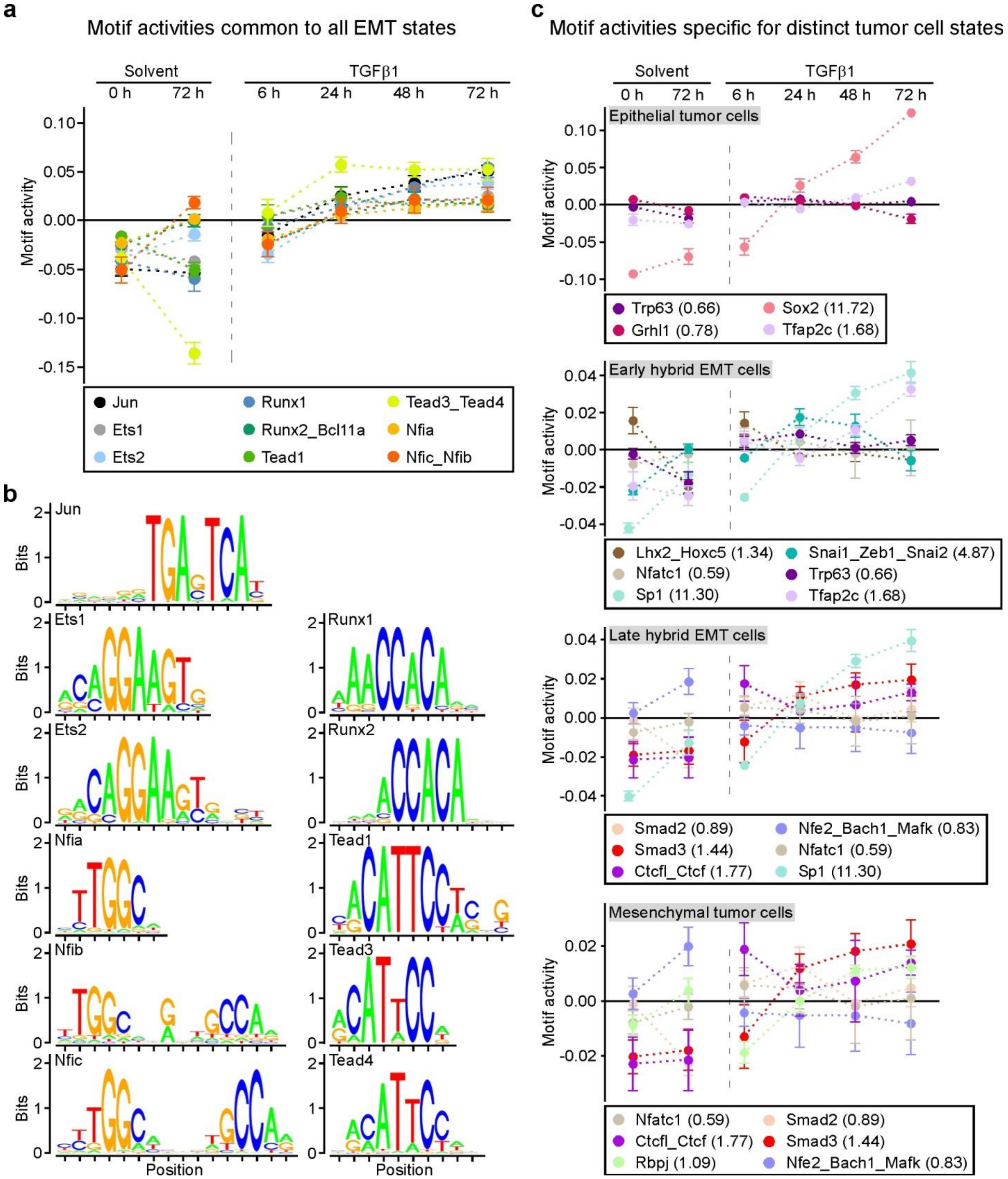


**Supplementary figure 8: Dynamics of transcription factor motif activity in the course of TGFβ1-induced partial EMT of oncogenically transformed organoids. a**, Activity plot of transcription factor motifs operating across the entire spectrum of purely epithelial, intermediate, and fully mesenchymal cells states (Pastushenko and Blanpain, 2019). ISMARA analyses were carried out with the RNA-seq data for TKA organoids (lines 931 and 947) harvested immediately at the onset of the experiment (0 h of cultivation; C-0), cultivated for 72 h in solvent (C-72), and treated with TGFβ1 for the indicated periods of time. Values shown represent the mean ± SD of the motif activities calculated from two independent biological replicates for both TKA organoid lines (931: n=2, 947: n=2). **b**, DNA sequence logos for the transcription factor motifs shown in (**a**). **c**, Activity plots for transcription factor motifs reported to be functional in epithelial, early hybrid EMT, late hybrid EMT, and mesenchymal tumor cell states (Pastushenko and Blanpain, 2019). Motif activities were determined and plotted as described in (**a**).

Citation: Pastushenko, I., and Blanpain, C. (2019). EMT Transition States during Tumor Progression and Metastasis. Trends Cell Biol 29, 212-226.

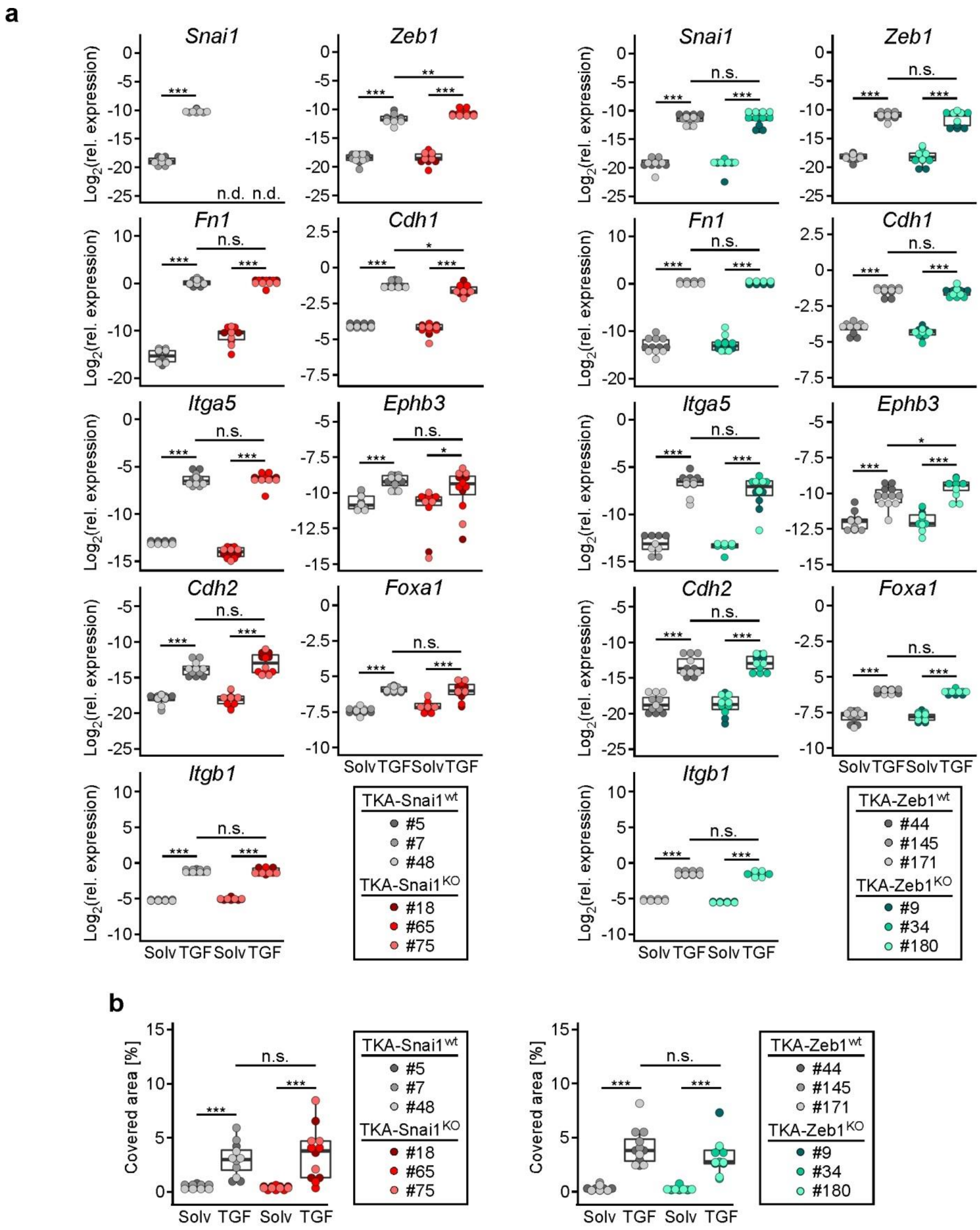


**Supplementary figure 9: Snail1 and Zeb1 are not required for the TGFβ1-induced partial EMT of oncogenically transformed organoids. a**, Gene expression analysis of EMT-TFs and EMT-associated genes in TKA-Snai1$^{wt}$, TKA-Snai1$^{KO}$, TKA-Zeb1$^{wt}$, and TKA-Zeb1$^{KO}$ organoid lines stimulated with solvent (solv) or TGFβ1 (TGF) for 72 h. Gene-specific transcripts were quantified by qRT-PCR and normalized to transcript levels of *Eef1a1* (n=4). Each dot represents the result of a single measurement while dot color identifies the organoid lines; n.d.: not detectable. ****p*<0.001**,** ***p*<0.01, **p*<0.05, n.s.: not significant; statistical significance was analyzed using the Mann-Whitney *U* test. Exact *p*-values are provided in Supplementary table 1. **b**, Invasiveness of TKA-Snai1$^{wt}$, TKA-Snai1$^{KO}$, TKA-Zeb1$^{wt}$, and TKA-Zeb1$^{KO}$ organoid lines treated with solvent or TGFβ1 for 96 h was quantified using Boyden chamber assays. Each dot represents the result of a single invasion assay while dot color identifies the organoid lines (n=4). ****p*<0.0001, n.s., not significant: *p*=0.7508 (TGFβ1-treated TKA-Snai1$^{WT}$ versus TKA-Snai1$^{KO}$) and *p*=0.1939 (TGFβ1-treated TKA-Zeb1$^{WT}$ versus TKA-Zeb1$^{KO}$); statistical significance was analyzed using the Mann-Whitney *U* test.

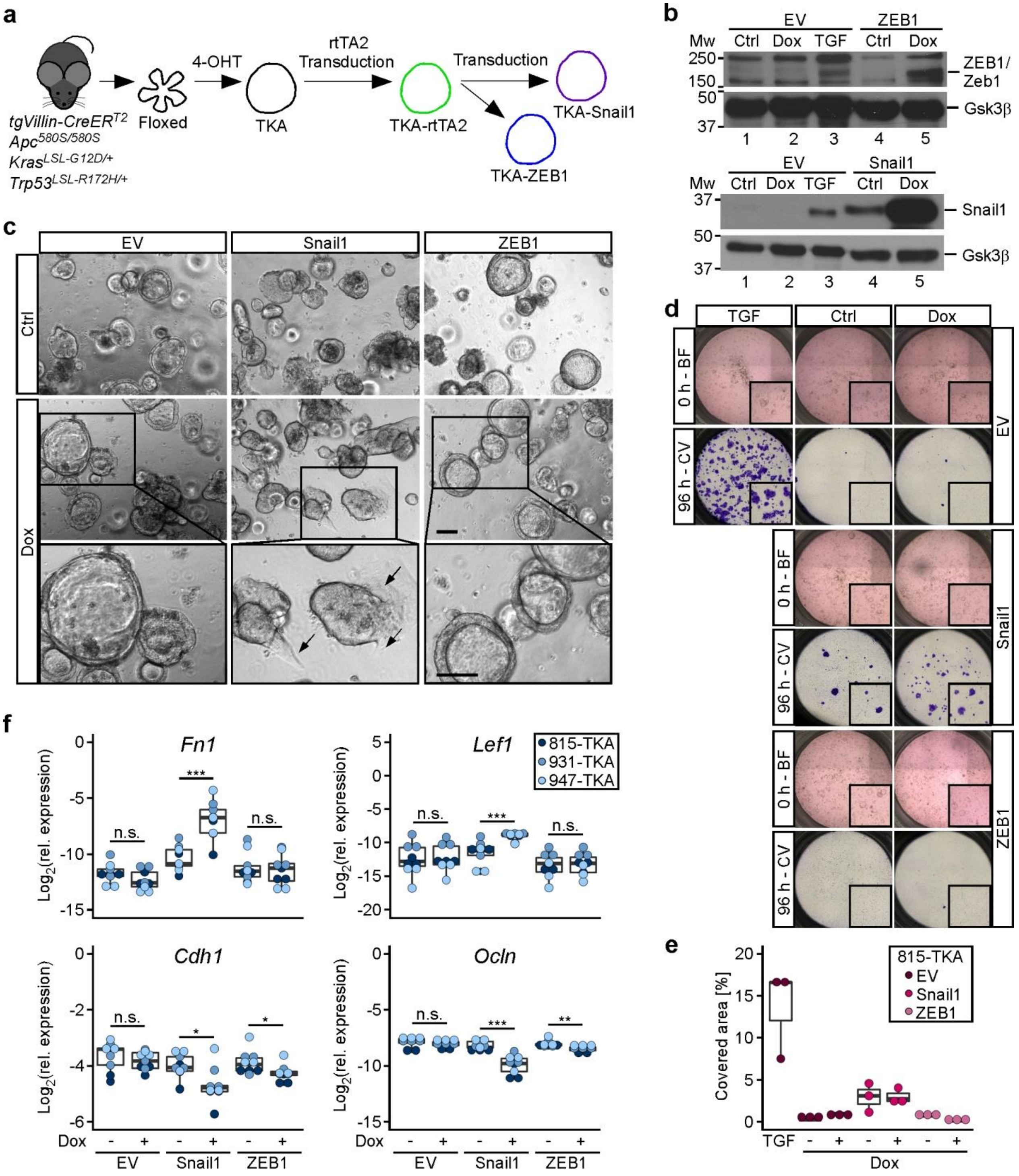


**Supplementary figure 10: Overexpression of Snail1 and ZEB1 does not mimic the TGFβ1-induced phenotype of oncogenically transformed organoids. a**, Strategy for the generation of TKA organoids with doxycycline (Dox)-inducible overexpression of murine Snail1 and human ZEB1. TKA organoids were transduced with a retroviral vector expressing the reverse tetracycline-dependent trans-activator (rtTA2), followed by a second transduction with an empty expression vector and retroviral constructs for Dox-inducible overexpression of Snail1 and

ZEB1. **b**, Western blot analyses to examine Snail1 and ZEB1 transgene expression in TKA organoids (line 815) left untreated (ctrl) or stimulated with Dox for 72 h. Gsk3β detection served as loading control. Results are representative for two independent biological replicates performed with derivatives of three different TKA organoid lines (815: n=2; 931: n=2; 947: n=2). Molecular weights of size standards are given in kDa. **c**, Phase contrast microscopic pictures of transduced TKA organoids (line 815) left untreated (ctrl) or stimulated with Dox for 72 h. Boxed areas are shown at higher magnification below. Arrows highlight sites of invasive behavior. Scale bars: 100 µm. **d**, Boyden chamber invasion assays with TKA organoids (line 815) transduced with empty vector (EV) and Dox-inducible expression vectors for Snail1 and ZEB1. Bright field (BF) images taken at the onset of the experiment (0 h) of untreated organoids (ctrl) and organoids receiving Dox. Empty vector-transduced organoids were treated with TGFβ1 for comparison. Inserts show magnified views of the upper chambers. Invaded cells were visualized by crystal violet (CV) staining after 96 h of treatment. Inserts show magnified views of the bottom faces of the Boyden chambers. **e**, Quantification of invasion experiments as shown in (**d**) performed with TKA organoids derived from line 815 and stimulated with TGFβ1 or cultured in the absence (-) or presence (+) of Dox for 96 h (n=3). Dots represent results of individual experiments. Dot color identifies the organoid lines. Statistical analysis was not appropriate due to low sample size. **f**, Gene expression analysis of *Fn1*, *Lef1*, *Cdh1*, and *Ocln* in TKA organoids transduced with empty vector (EV) and Dox-inducible expression vectors for Snail1 and ZEB1. Organoids were cultured in the absence (-) or presence (+) of Dox for 72 h. Gene-specific transcripts were quantified by qRT-PCR and normalized to transcript levels of *Eef1a1*. Dots represent results of individual experiments while dot color identifies the organoid lines. Three independent biological replicates each were performed with derivatives of three different TKA organoid lines (815: n=3; 931: n=3; 947: n=3). ***$p<0.001$, **$p<0.01$, *$p<0.05$, n.s.: not significant; Mann-Whitney *U* test. Exact *p*-values are provided in Supplementary table 1.
