## Supplementary tables and movie for "Canonical TGFβ signaling induces collective invasion in colorectal carcinogenesis through a Snail1- and Zeb1-independent partial EMT": Legend for Supplementary movie_Flum et al.docx

**Supplementary movie 1: TGFβ1-induced collective invasion of oncogenically transformed small intestinal organoids.** Live imaging of small intestinal TKA organoids (line 815) treated with solvent or TGFβ1 for 72 h. Images were acquired at 2 h intervals for a total of 72 h

.
