## Supplementary tables and movie for "Canonical TGFβ signaling induces collective invasion in colorectal carcinogenesis through a Snail1- and Zeb1-independent partial EMT": Suppl table 5_Flum et al.docx

**Supplementary table 5: DNA sequences of sgRNA targets and oligonucleotides employed in genome editing experiments (5’-3’)**

| **sgRNA target sequences excluding PAM** | |
| --- | --- |
| Smad4-sgRNA1 | ACTAATACCTTGACACTCTA |
| Smad4-sgRNA2 | GTTTTCAGTGGCTATTGATT |
| Snai1-sgRNA1 | GGTAGTCAACTCCGCTCGCG |
| Snai1-sgRNA2 | GTGTGGGTTGAGCCCGGATA |
| Zeb1-sgRNA1 | GATCTAGGCCTGCCATTCAC |
| Zeb1-sgRNA2 | TTATGAGTTCAAACCCATAG |
| **PCR primers for genotyping** | |
| Primer S1 (Smad4 forward primer) | CCCTTCTCCCCACCCTGATA |
| Primer S2 (Smad4 reverse primer) | TGCCTATGTGCAACCTCAGG |
| Snai1 forward primer | AGACAGTTCCAGGAACCCCT |
| Snai1 reverse primer | GCTGAAGCCTTCCCTCACTT |
| Zeb1 forward primer | AAGCCATACGAATGCCCGAA |
| Zeb1 reverse primer | TCGGCGATCTTTGAGAGCTC |
| Apc-A1 forward primer | GTTCTGTATCATGGAAAGATAGGTGGTC |
| Apc-A2 reverse primer | GAGTACGGGGTCTCTGTCTCAGTGAA |
| Apc-A3 reverse primer | CACTCAAAACGCTTTTGAGGGTTG |
| Kras-K1 forward primer | GTCTTTCCCCAGCACAGTGC |
| Kras-K2 reverse primer | CTCTTGCCTACGCCACCAGCTC |
| Kras-K3 forward primer | AGCTAGCCACCATGGCTTGAGTAAGTCTGCA |
| Trp53-T1 forward primer | AGCCTTAGACATAACACACGAACT |
| Trp53-T2 reverse primer | CTTGGAGACATAGCCACACTG |
| Trp53-T3 forward primer | GCCACCATGGCTTGAGTAA |
