## Supplementary tables and movie for "Canonical TGFβ signaling induces collective invasion in colorectal carcinogenesis through a Snail1- and Zeb1-independent partial EMT": Suppl table 6_Flum et al.docx

**Supplementary table 6: Plasmids used in the study**

| **Plasmid**  ***(purpose)*** | **derived from^a^** | **Source(s) of parental vectors** |
| --- | --- | --- |
| psPAX2  *(lentiviral packaging)* | n.a. | a gift from Didier Trono: Addgene plasmid #12260; RRID:Addgene_12260 |
| pMD2.G  *(lentiviral packaging)* | n.a. | a gift from Didier Trono: Addgene plasmid #12259; RRID:Addgene_12259 |
| Hit60  *(retroviral packaging)* | n.a. | a gift from Ulrich Maurer, University of Freiburg (Soneoka et al., 1995) |
| pVSV‐G  *(retroviral packaging)* | n.a. | (ClonTech, 631530) |
| pMSCV-rtTA2-PGK-eGFP-F2A-NeoR  *(expression of rtTA2)* | pMSCV-rtTA2-IRES-Eco-Receptor-PGK-PuroR | a gift from Cornelius Miething, University of Freiburg |
| pRetroX-tight-Snail1-HA-PuroR  *(Dox-inducible expression of Snail1-HA)* | n.a. | (Rönsch et al., 2015) |
| pRetroX-tight-ZEB1-HA-PuroR  *(Dox-inducible expression of ZEB1-HA)* | n.a. | (Schnappauf et al., 2016) |
| pRetroX-tight-MCS-PuroR  *(empty control vector)* | n.a. | (Rönsch et al., 2015) |
| pMSCV-loxP-BlastR-loxP-TGFBR1(T204D)-F2A-eGFP  *(4-OHT-inducible expression of TGFBR1(T204D)* | pcDNA3-ALK5 T204D  pMSCV-loxP-dsRed-loxP-eGFP-Puro-WPRE5 | a gift from Aristidis Moustakas; Addgene plasmid #80877; RRID:Addgene_80877)  a gift from Hans Clevers; Addgene plasmid #32702; RRID:Addgene_32702 |
| pMSCV-loxP-BlastR-loxP-TGFBR2(Δcyt)-F2A-eGFP  *(4-OHT-inducible expression of TGFBR2(Δcyt)* | pCMV5 HA-TBRII(delta Cyt)  pMSCV-loxP-dsRed-loxP-eGFP-Puro-WPRE5 | a gift from Joan Massague; Addgene plasmid #14051; RRID:Addgene_14051  a gift from Hans Clevers; Addgene plasmid #32702; RRID:Addgene_32702 |
| pMSCV-loxP-BlastR-loxP-eGFP  *(empty control vector)* | pMSCV-loxP-dsRed-loxP-eGFP-Puro-WPRE5 | a gift from Hans Clevers; Addgene plasmid #32702; RRID:Addgene_32702 |
| pLenti-SV40-mTomato-P2A-H2B-GFP-PuroR  *(expression vector for fluorescent proteins for life imaging)* | pcDNA6-mTomato-P2A-H2BGFP  pLenti-SV40-PuroR | a gift from Sebastian Arnold, University of Freiburg  this vector was generated based on pLenti CMV V5-LUC Blast, a gift from Eric Campeau; Addgene plasmid # 21474; RRID:Addgene_21474 |
| pCAG-Cas9-turbo-RFP plasmid  *(Cas9 expression vector)* | n.a. | (Freihen et al., 2020) |
| pMuLE ENTR U6-Smad4-sgRNA1-L1-R5  pMuLE ENTR U6-Snai1-sgRNA1-L1-R5  pMuLE ENTR U6-Zeb1-sgRNA1-L1-R5  *(sgRNA expression vectors)* | pMuLE ENTR U6 stuffer sgRNA scaffold L1-R5 | a gift from Ian Frew; Addgene plasmids # 62127; RRID:Addgene_62127 |
| pMuLE ENTR U6-Smad4-sgRNA2-L5-R4  *(sgRNA expression vector)* | pMuLE ENTR U6 stuffer sgRNA scaffold L5-L4 | a gift from Ian Frew; Addgene plasmids #62129; RRID:Addgene_62129 |
| pMuLE ENTR U6-Snai1-sgRNA2-L5-L2  pMuLE ENTR U6-Zeb1-sgRNA2-L5-L2  *(sgRNA expression vectors)* | pMuLE ENTR U6 stuffer sgRNA scaffold L5-L2 | a gift from Ian Frew; Addgene plasmids #62130; RRID:Addgene_62130 |
| pLenti‐Cas9‐T2A‐BlastR  *(viral Cas9 expression vector)* | n.a. | a gift from Jason Moffat; Addgene plasmid #73310; RRID:Addgene_73310 |
| pLenti-Dest-Snai1-sgRNA1+2-eGFP-F2A-NeoR-loxP  *(viral expression vector for Snail1-targeting sgRNAs)* | pLenti-Dest-eGFP  pMuLE ENTR U6-Snai1-sgRNA1-L1-R5  pMuLE ENTR U6-Snai1-sgRNA2-L5-L2 | a gift from Ian Frew; Addgene plasmid #62175; RRID:Addgene_62175 |
| pLenti-Dest-Zeb1-sgRNA1+2-eGFP-F2A-NeoR-loxP  *(viral expression vector for Zeb1-targeting sgRNAs)* | pLenti-Dest-eGFP  pMuLE ENTR U6-Zeb1-sgRNA1-L1-R5  pMuLE ENTR U6-Zeb1-sgRNA2-L5-L2 | a gift from Ian Frew; Addgene plasmid #62175; RRID:Addgene_62175 |

^a^: all plasmids were generated by standard cloning techniques; details are available upon request from the corresponding author.

n.a.: not applicable
