## Supplementary tables and movie for "Canonical TGFβ signaling induces collective invasion in colorectal carcinogenesis through a Snail1- and Zeb1-independent partial EMT": Suppl table 7_Flum et al.docx

**Supplementary table 7: List of antibodies used for immunofluorescence staining and Western blotting**

| **Primary antibodies for immunofluorescence staining** | | |
| --- | --- | --- |
| **Antigen** | **Type, origin, dilution** | **Supplier, catalogue number, clone number*** |
| β-catenin | polyclonal, rabbit, 1:50 | Cell Signaling Technology, #9581 |
| CD61 (integrin β3) | monoclonal, rabbit, 1:100 | Invitrogen, #MA5-32077, clone SJ19-09 |
| Claudin-7 | polyclonal, rabbit, 1:100 | Invitrogen, #34-9100 |
| Cleaved caspase-3 (Asp175) | polyclonal, rabbit, 1:400 | Cell Signaling Technology, #9661 |
| E-cadherin | monoclonal, mouse, 1:200 | BD Biosciences, #610182, clone 36/E-Cadherin |
| Fibronectin | polyclonal, rabbit, 1:400 | Abcam, #ab2413 |
| Ki67 | monoclonal, rabbit, 1:400 | Cell Signaling Technology, #9129, clone D3B5 |
| Laminin | polyclonal, rabbit, 1:25 | Sigma Aldrich, #L9393 |
| PKC ζ (atypical PKC isoform) | monoclonal, mouse, 1:100 | Santa Cruz, #sc-17781, clone H-1 |
| Smad2/3 | monoclonal, rabbit, 1:800 | Cell Signaling Technology, #8685, clone D7G7 |
| Vinculin | monoclonal, mouse, 1:400 | Sigma Aldrich, #V9131, clone hVIN-1 |
| **Secondary antibodies for immunofluorescence staining** | | |
| **Antigen** | **Origin, fluorophore, dilution** | **Supplier, catalogue number** |
| mouse IgG | donkey, Alexa Fluor555-conjugated, 1:500 | Invitrogen, #A-31570 |
| rabbit IgG | goat, Alexa Fluor488-conjugated, 1:200 | Invitrogen, #A-11008 |
| **Primary antibodies for Western blotting** | | |
| **Antigen** | **Type, origin, dilution** | **Supplier, catalogue number, clone number*** |
| α-tubulin | monoclonal, mouse, 1:10000 | Sigma Aldrich, #T9026, clone DM1A |
| Cleaved caspase-3 (Asp175) | polyclonal, rabbit, 1:1000 | Cell Signaling Technology, #9661 |
| E-cadherin | monoclonal, mouse, 1:1000 | BD Biosciences, #610404 |
| Ephb3 | monoclonal, mouse, 1:5000 | Abnova, #H00002049-M01, clone 3F12 |
| Fibronectin | polyclonal, rabbit, 1:1000 | Abcam, #ab2413 |
| Gsk3β | monoclonal, mouse, 1:1000 | BD Biosciences, #610201, clone 7/GSK-3β |
| Itga5 | polyclonal, rabbit, 1:1000 | Cell Signaling Technology, #4705 |
| Itgb1 | monoclonal, rabbit, 1:1000 | Cell Signaling Technology, #34971, clone D6S1W |
| Phospho-Smad2 (Ser465/467)/Smad3 (Ser423/425) | monoclonal, rabbit, 1:1000 | Cell Signaling Technology, #8828, clone D27F4 |
| Smad2/3 | monoclonal, rabbit, 1:5000 | Cell Signaling Technology, #8685, clone D7G7 |
| Smad4 | monoclonal, mouse, 1:2000 | Santa Cruz, #sc-7966, clone D7G7 |
| Snail1 | monoclonal, rabbit, 1:1000 | Cell Signaling Technology, #3879, clone C15D3 |
| Zeb1 | polyclonal, rabbit, 1:1000 | Sigma Aldrich, #HPA027524 |
| Zeb1^#^ | monoclonal, rabbit, 1:1000 | Cell Signaling Technology, XP®, #70512, clone E2G6Y |

*: if available

^#^: This antibody was applied only in experiments involving TKA organoids subjected to CRISPR/Cas9-mediated editing of the *Zeb1* gene
