## Supplementary tables and movie for "Canonical TGFβ signaling induces collective invasion in colorectal carcinogenesis through a Snail1- and Zeb1-independent partial EMT": Suppl table 8_Flum et al.docx

**Supplementary table 8: List of primer sequences used for qRT-PCR**

| **Gene** | **forward (5´ - 3´)** | **reverse (5´ - 3´)** |
| --- | --- | --- |
| Ascl2 | GGCTGTTAACACCCGCTACT | CTTTCCTCCGACGAGTAGGC |
| Axin2 | TCCCCACCTTGAATGAAGAA | TGGTGGCTGGTGCAAAGA |
| Cdh1 | CTTTAAGCCCAGCACTCAGG | CCTGCTTCCTGAGAAAATGC |
| Cdh2 | GGGAATCAGACGGCTAGACG | TCAGCAGCTTTAAGGCCCTC |
| Cdx2 | GAGTCCTGTGACCTCCTTGC | GGATTCTCGCAGCGTCCATA |
| Chga | GCCCGAAGTGACTTTGAGGA | CTACTCGAGCAGCAGTCTGG |
| Eef1a1 | GACAGCAAAAACGACCCACC | GGGCCATCTTCCAGCTTCTT |
| Ephb2 | CCAGCTTTAACACGGTGGAT | CCAGCTAGAGTGACCCCAAC |
| Ephb3 | GGTTTGCATCCTTTGACCTG | CTCGTTGGAGCTGAGTGTCA |
| Fn1 | TGACGCTGGCTTTAAGCTCA | TCATCCGCTGGCCATTTTCT |
| Foxa1 | AACCTCATGTCCTCCTCCGA | GCACGGGTCTGGAATACACA |
| Gapdh | ACCACAGTCCATGCCATCACT | GTCCACCACCCTGTTGCTGTA |
| Itga5 | ATTTCCGAGTCTGGGCCAAG | GATCCACAACGGGACACCAT |
| Itgb1 | GCGTGGTTGCTGGAATTGTT | AGGATTTTCACCCGTGTCCC |
| Krt20 | TAAAGACCCGGCTTGAGCAG | TTCAGAGGACACGACCTTGC |
| Lef1 | CAAGCGCCGACTTCCAAAAA | GAAGATGCTGGAGGATCGCA |
| Lgr5 | GGAAGACCTGAAGGCCCTTC | TCTGAACACGGTCAAAGCCA |
| Lyz | AGAGGGTGGTGAGAGATCCC | GGGAAAGCGAGGAAGTGTGA |
| Muc2 | GGATCACAGGTGCTCTTGCT | GCACAGACAGCTCTCGATGT |
| Ocln | CCTCCACCCCCATCTGACTA | GGAATCTCCTGGGCCACTTC |
| Snai1 | CTTGTGTCTGCACGACCTGT | CTTCACATCCGAGTGGGTTT |
| Zeb1 | GGGGCATCTCACACTTTTGT | AACGGCTGTGAACCAAAAAC |
